# Toxin-mediated downregulation of absorptive ion transporters NHE3, DRA, and SGLT1 in the colon contributes to diarrhea associated with *Clostridioides difficile* infection

**DOI:** 10.1101/2022.11.12.516162

**Authors:** F. Christopher Peritore-Galve, Izumi Kaji, Anna Smith, Lauren M. Walker, John A. Shupe, Pradeep K. Dudeja, James R. Goldenring, D. Borden Lacy

**Affiliations:** Department of Pathology, Microbiology, and Immunology, Vanderbilt University Medical Center, Nashville, TN, USA; Vanderbilt Institute for Infection, Immunology, and Inflammation, Vanderbilt University Medical Center, Nashville, TN, USA; Section of Surgical Sciences, Vanderbilt University Medical Center, Nashville, TN, USA; Epithelial Biology Center, Vanderbilt University School of Medicine, Nashville, TN, USA; Vanderbilt Vaccine Center, Vanderbilt University Medical Center, Nashville, TN, USA; Division of Gastroenterology and Hepatology, Department of Medicine, University of Illinois at Chicago, Chicago, Illinois, USA; Jesse Brown Veterans Affairs Medical Center, Chicago, Illinois, USA; Cell and Developmental Biology, Vanderbilt University School of Medicine, Nashville, TN, USA; Department of Veterans Affairs, Tennessee Valley Healthcare System, Nashville, TN, USA

**Keywords:** *Clostridioides difficile* infection, paracellular permeability, ion transport, toxins

## Abstract

**Background & Aim:** *Clostridioides difficile* infection (CDI) is the leading cause of hospital-acquired diarrhea and pseudomembranous colitis. Two protein toxins, TcdA and TcdB, produced by *C. difficile* are the major determinants of disease. However, the physiological cause of diarrhea associated with CDI is not well understood. We investigated the effects of CDI on paracellular permeability and apical ion transporters.

**Methods:** We studied intestinal permeability and apical membrane transporters in female C57BL/6J mice. Üssing chambers were used to measure regional differences in paracellular permeability and ion transporter function in intestinal mucosa. Intestinal tissues were collected from mice and analyzed by immunofluorescence microscopy and RNA-sequencing.

**Results:** CDI increased intestinal permeability through the size-selective leak pathway *in vivo*, but permeability was not increased at the sites of pathological damage. Chloride secretion was reduced in the cecum during infection by decreased CaCC function. Infected mice had decreased SGLT1 (also called SLC5A1) activity in the cecum and colon along with diminished apical abundance and an increase in luminal glucose. SGLT1 and DRA (also called SLC26A3) expression was ablated by either TcdA or TcdB, but NHE3 (also called SLC9A3) was decreased in a TcdB-dependent manner. Finally, expression of these three ion transporters was drastically reduced at the transcriptional level.

**Conclusions:** CDI increases intestinal permeability and decreases apical abundance of NHE3, SGLT1, and DRA. This combination may cause a dysfunction in water and solute absorption in the lower gastrointestinal tract, leading to osmotic diarrhea. These findings may open novel pathways for attenuating CDI-associated diarrhea.

**GRAPHICAL ABSTRACT:** 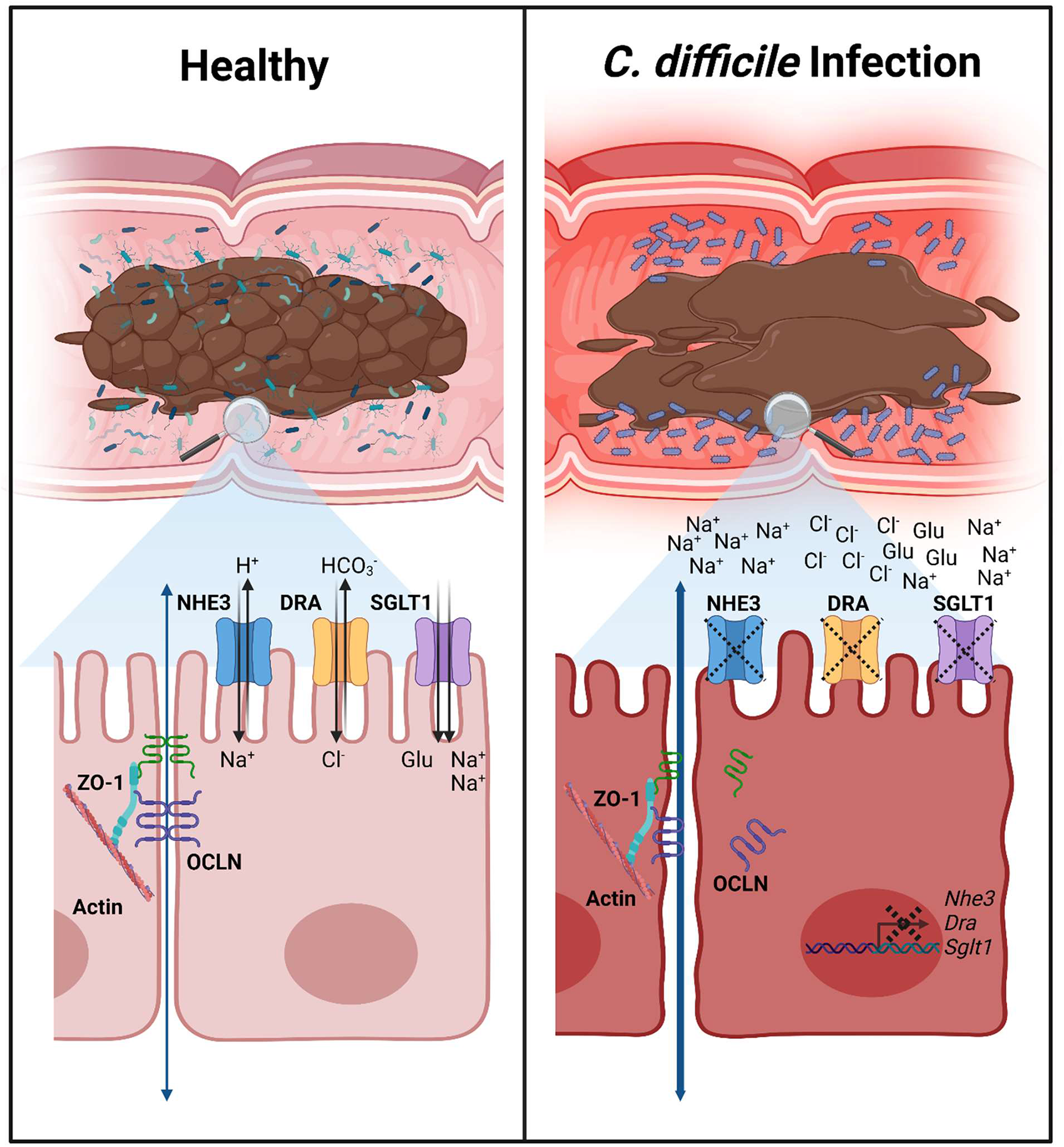

## INTRODUCTION

*Clostridioides difficile* infection (CDI) is the leading cause of hospital-acquired diarrhea in the USA, posing a significant burden on patients and the healthcare system^1, 2^. This antibiotic-associated infection causes approximately 250,000 cases and 14,000 deaths in the USA each year, which contributes $1.5 billion annually to the cost of healthcare^3, 4^. Furthermore, 25% of CDI patients experience a recurrent infection within two to eight weeks of the initial infection, with each episode increasing subsequent risk of recurrence^1, 5^. *C. difficile* is transmitted via the fecal-oral route by which hardy spores traverse the gastrointestinal (GI) tract to germinate in the ileum and colonize the colon as vegetative cells^6^. Infection results in a range of symptoms from mild to severe diarrhea, pseudomembranous colitis, and, in severe cases, toxic megacolon, sepsis, and death^7, 8^. Diarrhea provides a route of spore dispersal onto skin, bedding, clothes, and environmental surfaces that remain contaminated after symptom resolution^9, 10^. Environmental contamination is difficult to eliminate and perpetuates pathogen spread in both healthcare and community settings^10^. Therefore, a better understanding of the mechanisms underlying diarrhea during CDI and this symptom’s impact on disease severity may help improve patient outcomes and mitigate pathogen spread.

Pathogenesis during CDI is mediated by two large protein toxins, TcdA and TcdB, and may be exacerbated by a third toxin, CDT (*C. difficile* transferase)^11^. Upon production in the colon, TcdA and TcdB bind host cell receptors and are endocytosed^11–19^. Acidification of the endosome causes conformational changes in the toxins that lead to pore formation and delivery of the glucosyltransferase enzymes into the host cytosol. There, the enzymes irreversibly inactivate Rho-family GTPases by glucosylation, which disrupts the host cytoskeleton and leads to the production of proinflammatory cytokines, ultimately resulting in cytopathy and cell death^20–24^.

Additionally, TcdB concentrations at and above 0.1 nM induce glucosyltransferase-independent epithelial injury through the production of reactive oxygen species that results in a necrotic cell death^25–28^. Despite the understanding that TcdA and TcdB are the main determinants of disease, the role of each toxin during infection is not fully defined^28–32^. A recent study from our lab reaffirmed that TcdA and TcdB both cause symptoms in the murine model of infection and suggested that they synergize to worsen disease outcomes, including the severity of diarrhea^28^.

Early studies of *C. difficile* toxins revealed that injection of TcdA into rabbit ileal and colonic loops led to tissue damage, increased mucosal permeability, and fluid accumulation^33–35^. A follow up study demonstrated that TcdA, but not TcdB, caused up to a five-fold increase in intestinal permeability in the rabbit ileal loop model, suggesting that TcdA-induced paracellular permeability was a major factor in causing diarrhea^36^. Indeed, enhanced paracellular sodium and water efflux is an innate immune mechanism to clear *Citrobacter rodentium* colonic infections^37^. The idea that toxin-mediated tissue damage increases water and solute efflux through paracellular routes has been widely adopted as a likely mechanism of diarrhea in the *C. difficile* field. However, our understanding of diarrhea mechanisms have been improved based on the molecular identification of solute and water transporters during the past two decades. An *in vitro* study demonstrated that TcdB caused internalization of the Na^+^/H^+^ Exchanger 3 protein, NHE3 (a.k.a SLC9A3)^38^. Depletion of NHE3 by *C. difficile* was confirmed in CDI patient biopsies, which was accompanied by elevated Na^+^ and alkaline levels in CDI patient stool^39^. A more recent study demonstrated that the Cl^-^/HCO_3_^-^ exchanger protein DRA (Downregulated in Adenoma; a.k.a. SLC26A3) was depleted in CDI patient biopsies and during murine rectal instillation of TcdA, but not TcdB^40^. The coupled function of NHE3 and DRA is the primary route of electroneutral NaCl absorption in the lower GI tract, and functionally inactivating mutations in either gene cause congenital diarrhea^41–44^. Moreover, in models of inflammatory diarrhea, defects in paracellular permeability were not sufficient to cause diarrhea; rather, coordination of barrier dysfunction with impairment of NHE3 is necessary for solute malabsorption and diarrhea^45^. Together, these studies led us to hypothesize that CDI may induce concerted changes in paracellular permeability, ion absorption, and ion secretion that leads to spore-dispersing diarrhea. In this study, we examined this hypothesis with the aim of defining the mechanistic etiology of diarrhea during murine *C. difficile* infection.

## MATERIALS & METHODS

### Animals and Ethics Statement

Nine-week-old female C57BL/6J mice were purchased from The Jackson Laboratory and allowed to acclimate to the facility for one week before any procedures were performed. Mice were housed in specific pathogen-free conditions with 12-hour cycles of light and dark and used at 9-11 weeks of age. Cages were changed every two weeks to ensure clean bedding, and unless otherwise mentioned, they had free access to food and water. All studies were approved by the Animal Care and Use Committee at Vanderbilt University Medical Center and performed using protocol M1700185-01.

### Clostridioides difficile *Culturing*

Mutant and wildtype *C. difficile* strains (Table 1) were cultured at 37 °C in supplemented brain-heart infusion medium (BHIS) in an anaerobic chamber (90% N_2_, 5% H_2_, 5% CO_2_). Strains were stored at -80 °C in 20% glycerol stocks for long-term use. Spore stocks were prepared as previously described and stored at 4 °C until use^28, 46^.

### Clostridioides difficile *Infection*

Ten-week-old C57BL/6J mice were treated with cefoperazone (0.5 mg/ml) antibiotics in drinking water *ad libitum* for five days, then returned to normal drinking water two days before inoculation. Mice were inoculated with 10^5^ CFU/ml spores suspended in 100 µl sterile PBS by transoral gastric gavage. Spore stocks and diluted inoculum from mutant and wildtype *C. difficile* strains were serially diluted onto BHIS + taurocholic acid (TA; 10% w/v) media to enumerate concentration before and after inoculation.

Mice were monitored daily, and weight and symptom severity were recorded. Mouse stool samples were collected by scruffing mice and catching fresh stool in sterile, pre-weighed 1.5 ml microcentrifuge tubes. Stool samples were scored using a modified Bristol Stool Chart 1-4 scale by color and consistency. In this scoring scheme 1 = normal stool, 2 = well-formed but discolored stool, 3 = moist, soft, and discolored stool, and 4 = wet tail, watery diarrhea, and empty bowels^28^. Stool was then weighed, homogenized in 500 µl sterile PBS pH 7.4, and serially diluted on TCCFA media (TA 10% w/v; D-cycloserine 10 mg/ml; cefoxitin 10 mg/ml; fructose) to quantify *C. difficile* shed during infection. At the end of the experiment, mice were humanely euthanized by CO_2_ followed by cervical dislocation. The ilea, ceca, and colons were excised and imaged. Tissues were processed as described below based on the experiment performed. Weight change was plotted using GraphPad Prism and analyzed by two-way ANOVA with the Geisser-Greenhouse correction, then significant differences in weight loss were analyzed between wildtype and mutants using Tukey’s multiple comparisons test (*p* < 0.05). Stool scores presented were from day 2 for R20291 strains and vehicle control. Stool score was plotted using GraphPad Prism and analyzed using one-way ANOVA and unpaired t-test comparisons between wildtype and mutant strains (*p* < 0.05).

### *In vivo* Paracellular Permeability Measurement

Mice were pre-treated with antibiotics then inoculated with wildtype *C. difficile* R20291 (*n =* 5) or PBS (*n =* 6) as described above. Paracellular permeability was measured as described^47^. Two days post-inoculation, mice were placed into a new cage with water, but without food or bedding for 3 hours to clear gut contents. A probe solution containing 80 mg/ml fluorescein isothiocyanate (FITC) dextran (molecular weight 4 kDa; FD4) and 40 mg/ml rhodamine B dextran (molecular weight 70 kDa; RD70) was prepared in sterile milliQ water, then filter sterilized through a 0.2 µm filter into a new sterile tube. After three hours of fasting, 250 µl of probe solution was orally gavaged into each mouse, while noting the exact time of gavage. One of the mock-inoculated mice was orally gavaged with water to provide a control for plasma autofluorescence. Mice were returned to their food- and bedding-less cage and monitored. Food and bedding were returned 90 minutes post-gavage. Three hours post-gavage, mice were euthanized as described above, and blood was collected via heart stick using a 28G insulin syringe. Equal volumes of blood were placed into 1.3 ml collection tubes containing 3.2% sodium citrate (Sarstedt). Tubes were gently mixed by inversion to prevent coagulation. After all samples were collected, tubes were centrifuged at 1500 x g for 10 min at room temperature, and plasma was transferred to Spin-X UF 10 kDa molecular weight cutoff columns (Corning). These tubes were centrifuged for 10 min at room temperature to deproteinate the samples. Flowthrough was transferred into fresh microcentrifuge tubes. A standard curve was prepared, plasma fluorescein and rhodamine B fluorescence was measured on a BioTek Cytation 5 plate reader, and probe concentration in plasma was calculated exactly as described^47^.

### *Ex vivo* Paracellular Permeability Measurement in Üssing Chambers

To delineate the permeability of functionally distinct portions of the GI tract, mucosal-submucosal preparations of the ileum, cecum, proximal colon, and distal colon were collected from *C. difficile* R20291-infected and vehicle control mice at 2 days post-inoculation. Mice were euthanized, and tissues were excised by carefully cutting mesentery and connective tissue to prevent damage to the mucosa. Tissues were flushed with ice cold, oxygenated Krebs-Ringer Buffer (KRB) consisting of 117 mM NaCl, 25 mM NaHCO_3_, 11 mM glucose, 4.7 mM KCl, 1.2 mM MgCl_2_, 2.5 mM CaCl_2_, and 1.2 mM NaH_2_PO_4_. The segments were opened along the mesenteric border, and the seromuscular layer was carefully removed using fine forceps under a dissecting microscope in cold KRB. Preparations were made from each segment, mounted in 0.1 cm^2^ sliders, and placed in Üssing chambers (Physiologic Instruments Inc., Reno, NV). The luminal and serosal surfaces were bathed in 4 ml KRB maintained at 37 °C using a water-recirculating heating apparatus. To prevent prostaglandin synthesis, 10 µM of indomethacin was added immediately post-mounting to the serosal bath. To maintain pH at 7.4, the bathing solution was bubbled with a carbogen gas mixture of 95% O_2_, 5% CO_2_.

Mucosa-submucosa tissue preparations were allowed to stabilize for 60 min prior to application of fluorescence-conjugated dextrans. FD4 and RD70 were used to measure permeability in infected and control tissue segments through the leak and unrestricted pathways^47^. FD4 and RD70 were added to the serosal bath at concentrations of 0.25 mM and 17.5 mM, respectively. The mucosal bathing solution was sampled in technical duplicate at 0 and 60 minutes after addition of FD4 and RD70, and the bath volume was maintained by adding fresh KRB. Fluorescence intensity of each 100 µl sample and standards were analyzed at excitation/emission of 495 nm/525 nm for FD4 and 555 nm/585 nm for RD70 on a BioTek Cytation 5 plate reader^47, 48^. Paracellular permeability was determined as the concentration of FD4 and RD70 that passed through the mucosal-submucosal preparations from serosal to mucosal baths as µg/ml/cm^2^.

### Ion Transporter Functional Assessment in Üssing Chambers

To assess function of electrogenic ion transporters, short-circuit current (*I*_sc_), tissue conductance (G_t_, mS/cm^2^), and transmucosal resistance (R_t_, Ω · cm^2^) were recorded under voltage-clamp conditions. G_t_ was measured every 5 seconds with a bipolar pulse of ±3 mV for 20 mS, and R_t_ was calculated as the inverse of G_t_. The baseline *I*_sc_ and R_t_ were recorded after the fluorescent dextran permeability measurements before inhibitors were applied.

Sodium-glucose cotransporter 1 (SGLT1) activity in the presence of 11 mM glucose was determined as 0.1 mM phlorizin sensitive *I*_SC_. ENaC activity was determined as 10 µM amiloride sensitive *I*_SC_. Carbachol and forskolin (10 µM each) were sequentially applied to the serosal bath to compare Ca^2+^ and cAMP-dependent Cl^-^ secretory responses. The contribution of CFTR and CaCC to stimulated Cl^-^ secretion was measured by administering the CFTR inhibitor (R)-BPO-27 (10 µM) followed by CaCC-A01 (10 µM) after addition of forskolin.

After the Üssing protocol, mounted tissues were fixed in 10% Neutral Buffered Formalin (NBF) overnight, then embedded in paraffin by the Tissue Pathology Shared Resource at Vanderbilt University Medical Center (VUMC). Sections were mounted onto slides and stained with hematoxylin & eosin (H&E) to validate the removal of the seromuscular layer and lack of ischemic cell death during the experiment. H&E-stained slides were imaged at 40x magnification using a Leica SCN400 Slide Scanner at the Digital Histology Shared Resource at VUMC.

### Immunofluorescence Staining, Imaging, and Analysis

Mice were euthanized two days post-inoculation, then the colon of each animal was excised. Colons were gently flushed with cold sterile PBS, then cut transversally and fixed in 2% paraformaldehyde (PFA) at 4 °C for 2 hours or 10% NBF at 4 °C for 16 hours. After two hours, PFA-fixed tissues were washed three times in cold sterile PBS, then placed in a cold solution of 30% sucrose/1% NaN_3_ and incubated for 16 hours at 4 °C. Tissues were then Swiss rolled and embedded in OCT (Optimal Cutting Temperature embedding medium; Fisher Healthcare) in dry ice-cooled ethanol and stored at -80 °C until use. NBF-fixed tissues were rinsed in cold sterile PBS, then Swiss rolled and embedded in paraffin wax by the TPSR.

Formalin-fixed paraffin embedded (FFPE) blocks were sectioned into 7 µm slices on a Microm HM 335E microtome, then dried overnight and kept at room temperature until use. Paraffin wax was removed using xylenes, then tissues were rehydrated in an ethanol gradient. Antigen retrieval was performed using 0.1 M Citrate Buffer pH 6.0 before the blocking step. Frozen colon blocks from mice inoculated with *C. difficile* strains or vehicle control were sectioned into 7 µm slices on a Leica CM1950 Cryostat and kept at -80 °C until use. OCT medium on frozen slides was gently removed by incubating in three sequential PBS washes. Both OCT and FFPE tissues were blocked for an hour at room temperature in a solution of PBS pH 7.4, 2% normal goat or donkey serum, and 0.3% Triton X-100.

Afterwards, the blocking solution was removed and replaced with the primary antibody diluted at a specified concentration (Table 1) in an antibody dilution buffer comprised of PBS, 1% bovine serum albumin (BSA), and 0.3% Triton X-100. Slides were incubated with the primary antibody overnight at 4°C. The following day, slides were washed in three changes of PBS, then incubated for an hour at room temperature in antibody dilution buffer containing secondary antibodies and phalloidin (Table 1). After an hour, slides were washed in three changes of PBS, incubated in DAPI for 1 minute, then washed in PBS. Slides were cover slipped with ProLong Gold and allowed to cure overnight before sealing with clear nail polish and placing at 4 °C in the dark.

Sections were imaged using a Nikon Spinning Disk confocal microscope equipped with a Yokogawa CSU-X1 spinning disk head and a Photometrics Prime 95B sCMOS monochrome camera, and 405 nm, 488 nm, 561 nm, and 647 nm diode laser lines. For each colon sample, at least three distinct regions of the proximal colon (first 1/3^rd^ of colon) and distal colon (last 1/3^rd^ of colon) were imaged for analysis in at least three mice per treatment. These images were acquired at 20x with a Plan Apo Lambda 20x 0.75 NA WD 1.00mm objective lens. Magnified images were acquired at 40x, 60x, or 100x with Pan Fluor 40x Oil DIC H N2 1.30 NA WD 0.20mm, Plan Apo Lambda Oil 60x 1.40 NA WD 0.13mm, and Apo TIRF Oil 100x 1.49 NA WD 0.12mm objective lenses, respectively.

Relative fluorescence intensity was quantified as a metric for protein abundance using FIJI software (National Institutes of Health, Bethesda, MD). A baseline threshold was generated with vehicle control images using Otsu’s method^49^, and fluorescence intensity by area was calculated. The same threshold settings were applied across all treatments and regions of the colon for each individual marker. Fluorescence intensity data was averaged between different fields of view from the same region of the same mouse, and relative fluorescence intensity was calculated with reference to the vehicle control.

### Stool Glucose Measurement

Stool was collected into sterile, pre-weighed microcentrifuge tubes from mice inoculated with *C. difficile* R20291 (*n =* 5 per treatment) at day 0, prior to inoculation, and two days post-inoculation. Tubes containing stool were weighed, and glucose was measured using the Glucose Assay Kit (Abcam) according to manufacturer’s instructions. Briefly, stool was mechanically homogenized in equal volumes of ice-cold assay buffer, then centrifuged at 10,000 x g for 5 min at 4 °C. The supernatant was transferred to Spin-X UF 10 kDa molecular weight cutoff columns (Corning) and centrifuged at 10,000 x g for 10 min at 4 °C to deproteinate the samples. Flowthrough was then used for colorimetric quantification of glucose concentration compared to standards provided by the manufacturer.

### Transcriptome Profiling of the Distal Colon

Mice inoculated with *C. difficile* wildtype and mutant strains, or the vehicle control were euthanized at 2 days post-inoculation, and a 0.5 cm segment of the distal colon was harvested and placed in RNAlater solution on ice. Tissues were washed with sterile PBS and then transferred to fresh tubes containing 1 ml of TRIzol and 1 mm diameter Zirconia-silica beads. Colon samples were homogenized in an Omni Bead Ruptor 4 homogenizer for one minute in two 30 s intervals, then placed on ice for five minutes. Phases were separated by addition of chloroform and centrifugation, then total RNA was precipitated, washed, and solubilized in nuclease-free water. Contaminating DNA was depleted using TURBO DNA-free Kit (Thermo Fisher), then RNA was cleaned using the Monarch RNA Cleanup Kit (NEB). Quantity and quality were assessed by spectroscopy.

RNA-sequencing libraries were prepared by depleting eukaryotic ribosomal RNA with the NEBNext rRNA Depletion Kit prior to library synthesis with the NEBNext Ultra II Directional RNA Library Prep Kit for Illumina and addition of multiplex oligos using the Unique Dual Index Primer Pairs Set 3 (NEB). Library quality control was performed using Qubit and Bioanalyzer 2100. Paired end 150 bp sequencing was performed by the VANTAGE Core on the Illumina NovaSeq 6000, with the aim of generating 50 million reads per sample. Sequencing data quality was assessed with FASTQC, then reads were aligned to the GRCm39 mouse genome (GenBank accession number GCA_000001635.9) using the STAR alignment tool with default settings. Normalization by trimmed mean of M-values and differential expression was performed with limma-voom. Plots of differentially expressed genes were generated using GraphPad Prism, or with Heatmapper^50^. Raw and analyzed transcriptomics data have been deposited on NCBI’s Gene Expression Omnibus and are accessible through GEO Series accession number GSE216919.

### Statistical Analyses

Data were plotted with GraphPad Prism, and statistical comparisons were determined using one-way ANOVA and unpaired Student’s t-test for two-group comparisons. For more than three groups, two-way ANOVA and Tukey’s multiple comparisons test was performed. *P* < 0.05 was considered as a minimal significant difference.

## RESULTS

### Both TcdA and TcdB contribute to diarrhea severity during murine CDI

We previously observed that diarrhea was most severe in mice infected with the wildtype *C. difficile* R20291 strain when compared to mutants that had mutationally impaired glucosyltransferase activity in either TcdA or TcdB (A+ B_GTX_ and A_GTX_ B+)^28^. To generate tissues for the experiments described below, we repeated previously published disease analyses and found consistent results^28^. Weight loss was recorded over the course of 2 days post-inoculation (dpi), revealing significantly reduced percentage weight loss at 2 dpi between R20291 and A+ B_GTX_ (17.1% avg. weight loss vs. 5.6%, *p* = 0.018), R20291 and A_GTX_ B+ (17.1% avg. weight loss vs. 7.1%, *p* = 0.0277), and R20291 and Δ*tcdA*Δ*tcdB* (17.1% avg. weight loss vs. 1.68% avg. weight gain, *p* < 0.0001) (Figure 1A). Mice inoculated with A+ B_GTX_ and A_GTX_ B+ strains lost significantly more weight than Δ*tcdA*Δ*tcdB*-inoculated mice (*p* < 0.05; contrast not shown on graph), but there was no difference in weight loss between A+ B_GTX_ and A_GTX_ B+ (Figure 1A). Stool samples were scored at 2 dpi, demonstrating that R20291 consistently caused more severe diarrhea compared to toxin mutant strains (*p* < 0.001; Figure 1B). These results are consistent with the empty ceca and colons associated with WT infected mice (Figure 1C). *In situ* assessment of the GI tract highlighted inflamed and empty bowels observed during infection with the R20291 strain (Figure 1D). These data indicate that both TcdA and TcdB contribute to weight loss and diarrhea in our mouse model of CDI and provide the framework for our decision to examine the role of toxins in R20291-associated diarrhea at 2 dpi.

**FIGURE 1.**
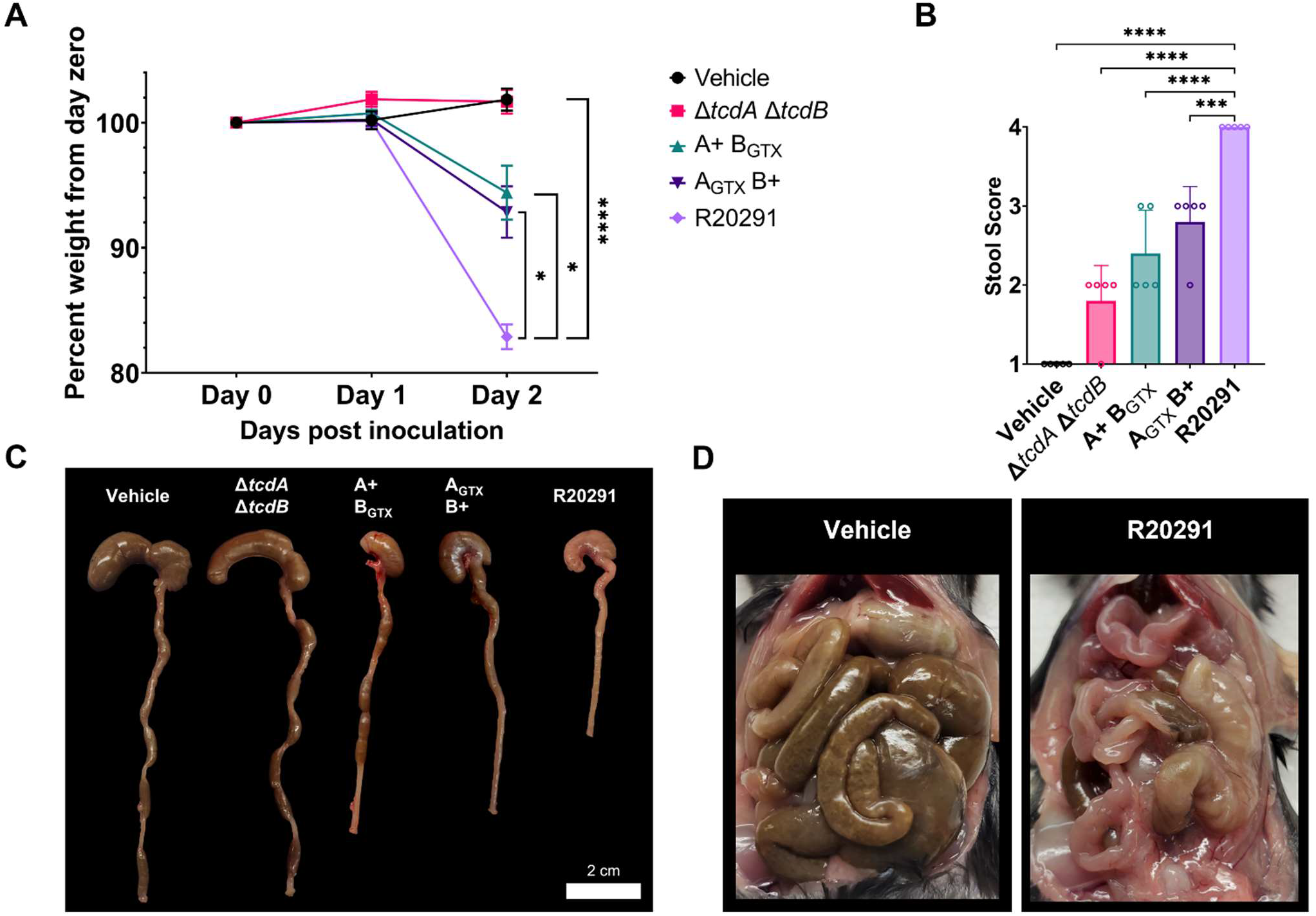
**A)** Percent weight loss of mice inoculated with wildtype and mutant *C. difficile* strains demonstrates the toxin-additive nature of weight loss. (*n =* 5 per strain or control) **B)** Stool scores are consistently more severe in wildtype R20291 compared to toxin mutant strains (*n* = 5 per strain or control). **C)** Representative images of mouse ceca and colons during CDI with wildtype and mutant strains at 2 dpi. Wildtype-inoculated mice have empty ceca and colons and appear inflamed. Scale bar = 2 cm. **D)** Abdominal cavities of representative mice inoculated with either R20291 or the vehicle control at 2 dpi. * *P* < 0.05; *** *P* < 0.001; **** *P* < 0.0001.

### Intestinal permeability is increased via the leak pathway

To define the pathways involved in *C. difficile* diarrhea, we first measured intestinal permeability in the murine model of infection using the wildtype R20291 strain. Oral gavage of FD4 and RD70 in fasted mice inoculated with R20291 or the vehicle control demonstrated that intestinal permeability is increased during CDI through the leak pathway (*p* = 0.038), but not the unrestricted pathway (Figure 2A). We predicted that FD4 permeability would be increased in the lower GI tract, where *C. difficile* germinates (ileum) and causes damage (cecum and colon). Tissue segments from the ileum, cecum, proximal colon, and distal colon were excised from R20291- and vehicle-inoculated animals at 2 dpi, then prepared and mounted in Üssing chambers. FD4 and RD70 were added to the serosal bath, and permeability was measured after 60 min. Unexpectedly, there was no increase in FD4 or RD70 permeability in R20291 infected mice (Figure 2B and Supplementary Figure 1). As an orthogonal measurement for barrier integrity, baseline transmucosal resistance (R_t_) was measured 60 minutes after mounting, which revealed no significant change in R_t_ between CDI and healthy mice in the regions tested (Figure 2C). After 60 min, these tissue segments were subjected to a course of ion transport inhibitors and stimulants to test electrogenic ion transport during CDI. Despite the exposure of chemicals over the course of 120 min, there were no visible effects of damage resulting from ischemia that would bias the results of barrier function and ion transport (Figure 2D). Together, these results suggest that CDI increases the paracellular leak pathway but not in the expected tissue areas tested within the Üssing chamber (i.e.. ileum, cecum, proximal colon, and distal colon). The Üssing chamber experiments also indicate that CDI has minimal effect on epithelial barrier function from the ileum to the distal colon despite the pathological damage caused during infection.

**FIGURE 2.**
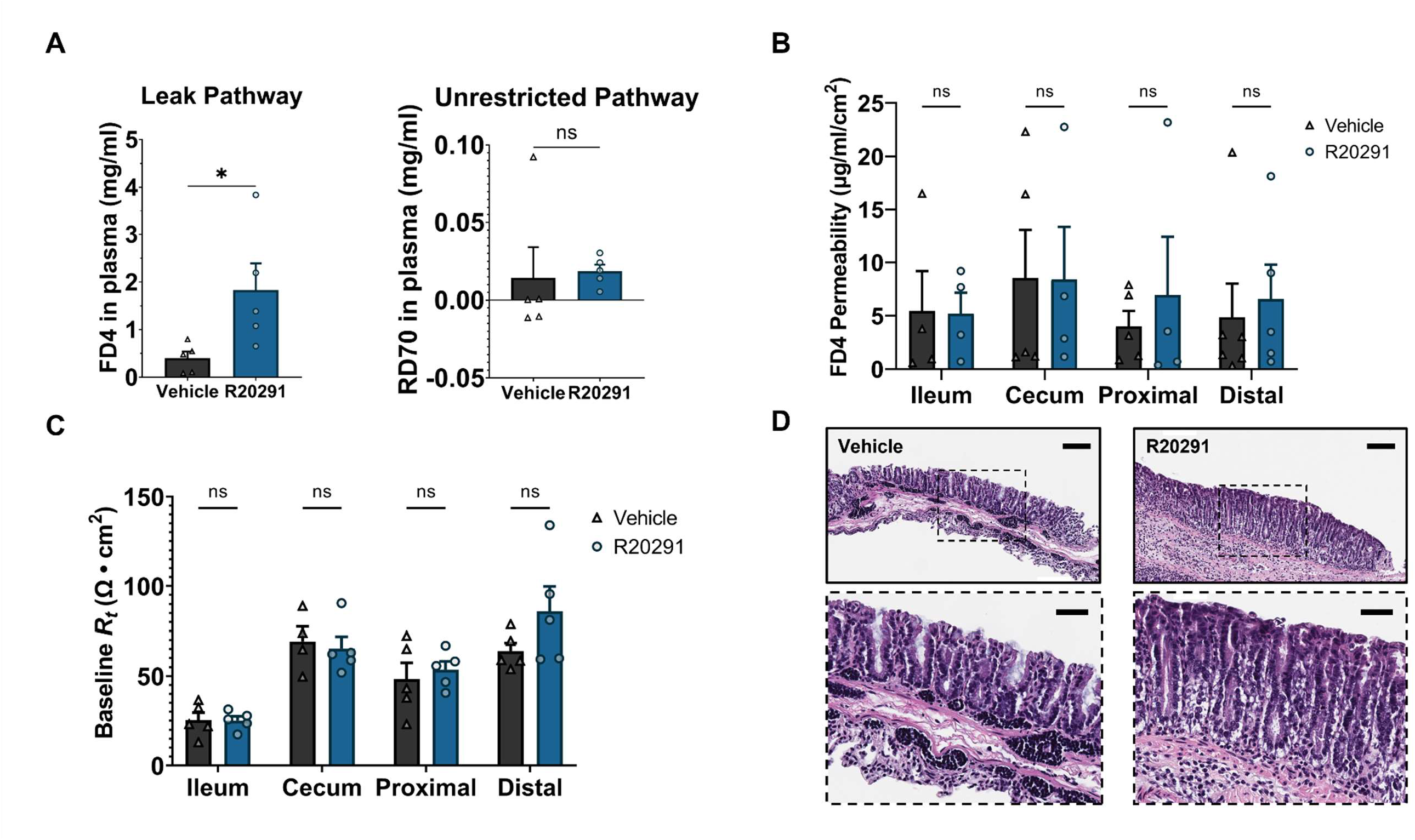
**A)** Paracellular flux is increased via the leak pathway during CDI, and there is no increase in permeability through the unrestricted pathway (*n =* 5 per treatment). **B)** There is no difference in paracellular permeability between mock- and R20291-inoculated mice at 2 dpi via the leak pathway in the ileum, cecum, proximal colon, and distal colon (*n* = 4-5 per treatment per region). **C)** Baseline transmucosal resistance is unaffected during CDI in the lower GI tract (*n* = 4-5 per treatment per region). **D)** Representative H&E-stained mucosal preparations of distal colons after an Üssing chamber experiment, exemplifying the lack of ischemia during the protocol. Scale bars = 100 µm. * *P* < 0.05.

### *C. difficile* infection causes only minor effects on electrogenic ion transport

The function of apical ion transporters in the lower GI tract during CDI were tested in Üssing chambers. The baseline short circuit current (*I*_sc_) was not significantly different between R20291 and control mice, suggesting a minimal effect of CDI on electrogenic ion transport (Figure 3A). The Epithelial Sodium Channel (ENaC), which is functionally relevant in the distal colon, was tested through inhibition with amiloride. There was no significant impairment of ENaC function between R20291 and control mice (*p* < 0.05) (Figure 3B and Supplementary Figure 2). Cystic Fibrosis Transmembrane Conductance Regulator (CFTR) is responsible for most anion secretion in response to increases in intracellular Ca^2+^ and cyclic dinucleotides, and Calcium activated Chloride Channels (CaCC) contributes to Cl^-^ secretion. To assess their function during CDI, the acetylcholine analogue, carbachol (Cch), and the adenylate cyclase-inducing chemical, forskolin (Fsk), were applied sequentially. Addition of each molecule revealed that the secretory potential of the cecum was nearly halved during CDI, while there was no effect on other regions of the lower GI tract (Figures 2C and 2D). The CFTR-dependent portion of stimulated secretion was measured by adding a maximal dose of the inhibitor R-BPO-27. This revealed that CFTR function was not significantly impaired during CDI in the lower GI tract (Figure 3E). CaCC-dependent secretion was measured by the addition of the inhibitor CaCC-A01. Activity of CaCC was reduced approximately five-fold in the cecum during CDI, accounting for the reduction in secretion potential when stimulated with Cch and Fsk (Figure 3F). CaCC function was not impaired in any other regions of the lower GI during CDI (Figure 3F). These results suggest that Cl^-^ hypersecretion is likely not a cause of diarrhea, nor is ENaC-mediated sodium malabsorption.

**FIGURE 3.**
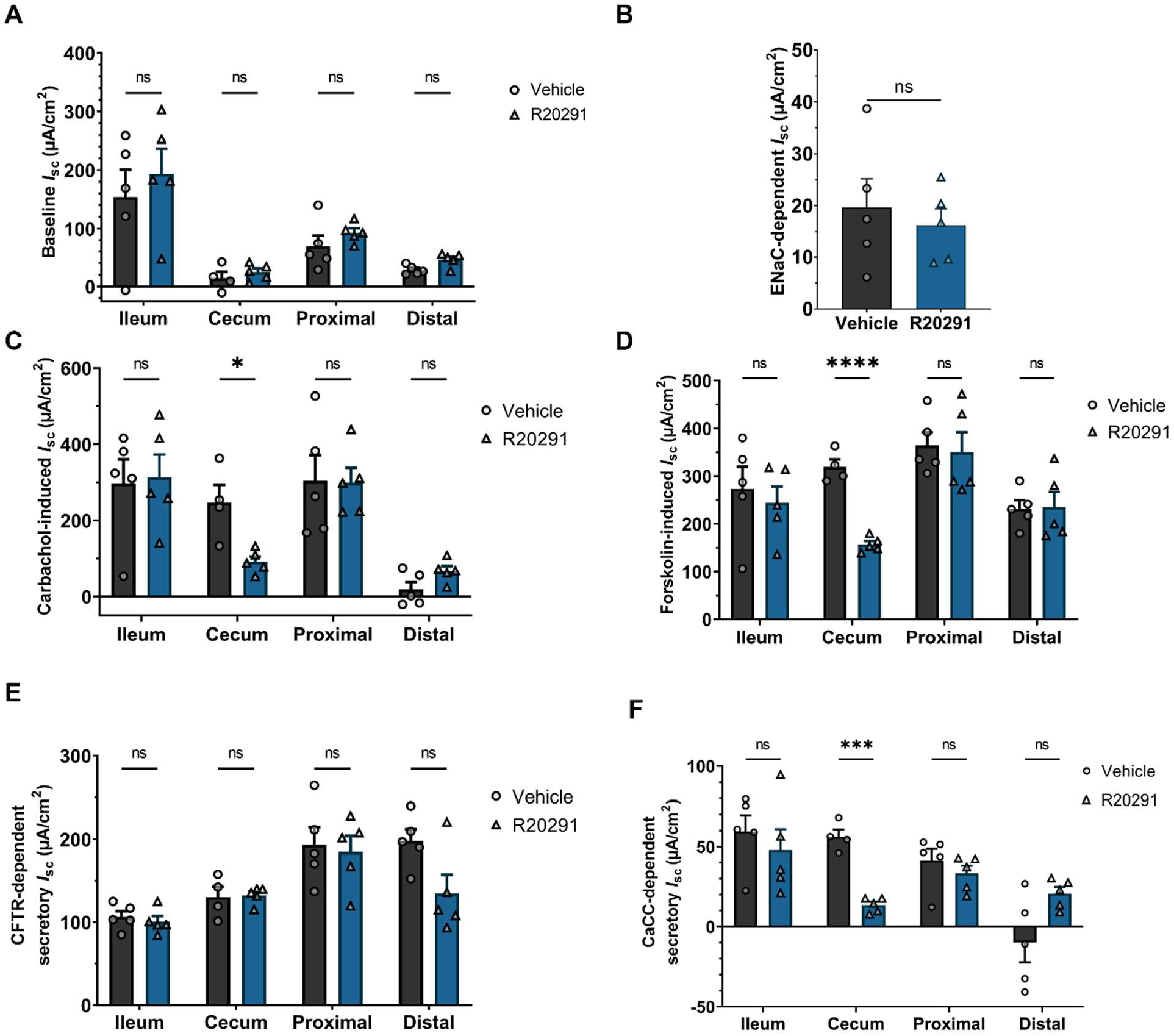
**A)** Baseline short circuit current (*I*_sc_) is unaffected during CDI in the lower GI tract (*n* = 4-5 per treatment per region). **B)** The function of the absorptive sodium channel, ENaC, is unchanged during CDI. **C)** The acetylcholine analogue carbachol induces Ca^2+^ induced Cl^-^ secretion, revealing the reduced Cl^-^ secretory potential in the cecum alone during CDI. **D)** The adenylyl cyclase activator forskolin induces cAMP-dependent Cl^-^ secretion, again revealing the reduced Cl^-^ secretory potential in the *C. difficile*-infected cecum. **E)** CDI causes no significant effect on Cystic Fibrosis Transmembrane Conductance Regulator (CFTR) function in the lower GI tract. **F)** Calcium activated Chloride Channel (CaCC) activity is reduced significantly in the cecum only during CDI. * *P* < 0.05; *** *P* < 0.001; **** *P* < 0.0001.

### *C. difficile* infection impairs SGLT1 function and expression in the colon

We also determined the function of sodium-glucose linked co-transporter (SGLT1; SLC5A1), which is critical for absorption of glucose/galactose, Na^+^, and water through the apical cell membrane in the small intestine^51, 52^. The function of SGLT1 in the presence of 11 mM glucose was tested through inhibition by phlorizin. There was no effect in SGLT1 function in the ileum, but unexpectedly, there was a significant reduction in function in the cecum, proximal colon, and distal colon during CDI (*p <* 0.05) (Figure 4A). Using immunofluorescence microscopy, we found that SGLT1 expression was completely ablated in the colon during infection with R20291 (Figure 4B). SGLT1 is responsible for the majority of glucose uptake, with most of it taking place in the small intestine. The physiological relevance of SGLT1 in the colon is incompletely understood, but analysis of fecal samples revealed that there is a two-fold increase in stool glucose in R20291-infected mice at the same timepoint at which expression of SGLT1 function and expression is decreased, suggesting that SGLT1 in the colon may be important in reabsorbing excess glucose (Figure 4C). Additionally, the role of SGLT1 in carbohydrate, sodium, and water absorption is consistent with a model where loss of SGLT1 expression could contribute to diarrhea.

**FIGURE 4.**
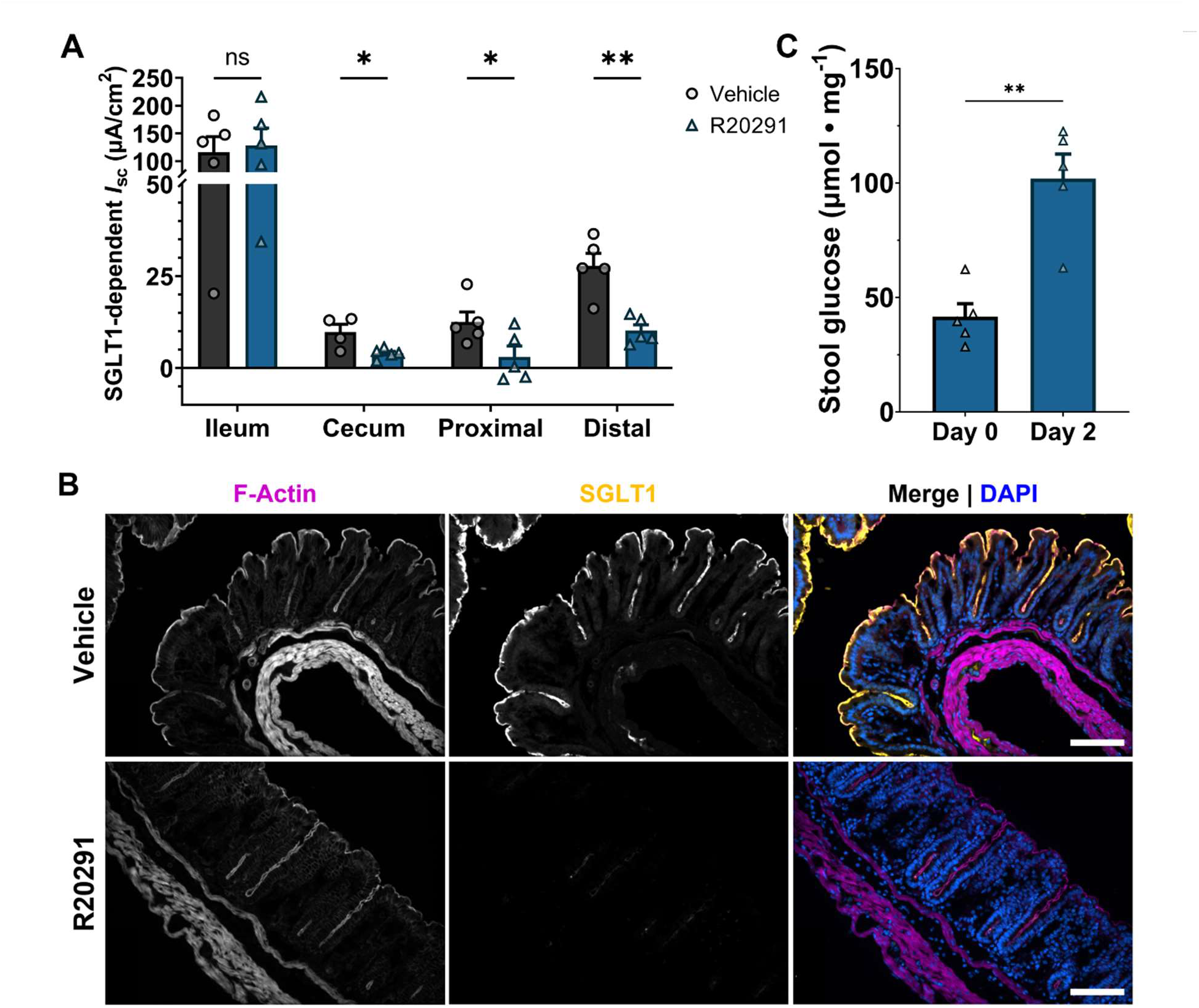
**A)** Sodium-glucose cotransporter 1 (SGLT1) activity is significantly reduced in the cecum, proximal colon, and distal colon during CDI. **B)** SGLT1 expression is completely ablated at the apical surface of the colonic mucosa during infection. Scale bars = 50 µm. **C)** Stool glucose concentration is increased two-fold during infection. * *P* < 0.05; ** *P* < 0.01; *** *P* < 0.001; **** *P* < 0.0001.

### TcdB is the major driver of ion transporter depletion

NHE3 and DRA are depleted during human CDI, thus are predicted factors in causing diarrhea^38–40^. Given our observation that both TcdA and TcdB contribute to weight loss and diarrhea (Figure 1C), we hypothesized that each toxin could have a different effect on expression of ion transporters implicated in diarrhea. Using immunofluorescence microscopy, we found that SGLT1 expression was significantly decreased in the distal colon in response to either or both toxins but was largely present during colonization with a toxin-negative strain (Figures 5A and 5B). Specifically, expression of SGLT1 was ablated in the presence of functional TcdB, but in the A+ B_GTX_-infected mice, partial expression remained in the crypt base (Figure 5A).

**FIGURE 5.**
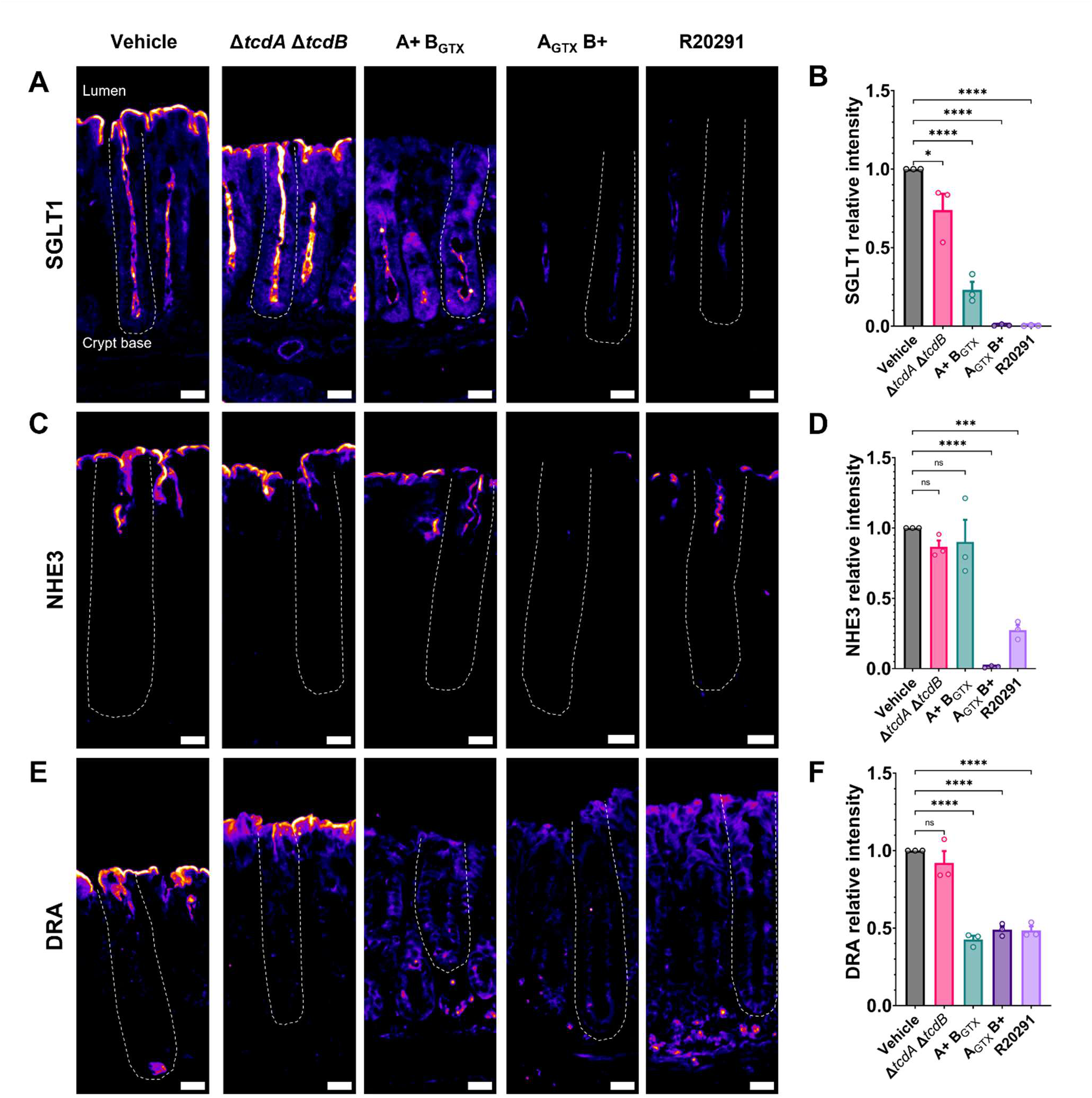
**A & B)** SGLT1 immunofluorescent expression is presented with Fire Lookup Table settings, where higher abundance correlates to more intense gold color. SGLT1 expression in the distal colon is decreased completely by TcdB, but low crypt expression remains in the presence of TcdA alone. SGLT1 expression is slightly decreased after inoculation with the double knockout strain, Δ*tcdA* Δ*tcdB*, but is decreased more drastically by toxigenic *C. difficile* strains. **C & D)** NHE3 distal colon expression is severely decreased only when TcdB is present and functional. **E & F)** DRA distal colon expression is decreased by either TcdA or TcdB. * *P* < 0.05; *** *P* < 0.001; **** *P* < 0.0001. Scale bars = 50 µm.

NHE3 is present in a gradient of high-to-low expression from the proximal to the distal colon in small animal models^53^. No studies have examined NHE3 expression in animal models of CDI. Unexpectedly, NHE3 expression in the proximal colon was not affected by infection with any *C. difficile* strain (Supplementary Figure 3). However, expression was significantly decreased in a TcdB-dependent manner in the distal colon (Figure 5C and 5D).

NHE3 is internalized in response to TcdB glucosyltransferase activity *in vitro*^38^. It was hypothesized that internalization could be caused by TcdB-dependent deactivation of Rho-GTPases, which in turn disrupts stabilization of the cytoskeletal protein group Ezrin/Radixin/Moesin (ERM), which is linked to Na^+^-H^+^ exchanger regulatory factor (NHERF) family proteins that anchor NHE3 to the cell membrane^38, 54–56^. We used immunofluorescence microscopy to quantify expression and observe localization of NHERF1 and the active, phosphorylated ERM (pERM). There was no significant effect on NHERF1 or pERM expression and localization during infection with R20291, suggesting the potential for another mechanism of NHE3 depletion during infection (Supplementary Figures 4 and 5).

DRA is functionally linked to NHE3 and is expressed in a low-to-high gradient from the proximal to distal colon in rodents^53^. DRA has previously been shown to be downregulated at the protein but not mRNA level in mice intrarectally instilled with TcdA, TcdA and TcdB, but not TcdB alone^40^. Immunofluorescence analysis revealed that DRA expression was significantly decreased in the distal colon in response to functional TcdA or TcdB, with no additive outcome when both functional toxins were present (Figures 5E and 5F). Together, these data demonstrate impact of TcdA and TcdB on the expression of absorptive ion transporters in the distal colon, transporters that may be major factors in driving diarrhea.

### Ion transporter transcription is decreased through toxin-dependent mechanisms

Since NHERF1 and pERM expression were not altered during CDI, we hypothesized that NHE3 was decreased at the transcriptional level. We used whole genome RNA-sequencing as an unbiased approach to uncover toxin-dependent transcriptional effects. Transcriptomic analysis of the distal colon identified the potent upregulation of genes involved in the innate immune response including *Retnlb, Clca4b, Prss22, S100a9, Chil3,* and *Saa3* (Figure 6A and Supplementary Table 1). The most highly downregulated gene during infection was the cytochrome p450 family protein-encoding gene *Cyp2c55*, which has been previously reported^31^ (Figure 6A). Of relevance to this study, the second most highly downregulated gene was *Dra* (S*lc26a3*), followed by other solute and water transporters *Aqp8*, *Clca1*, *Slc20a1*, *Slc26a2*, *Nhe3* (*Slc9a3*), and *Sglt1* (*Slc5a1*).

**FIGURE 6.**
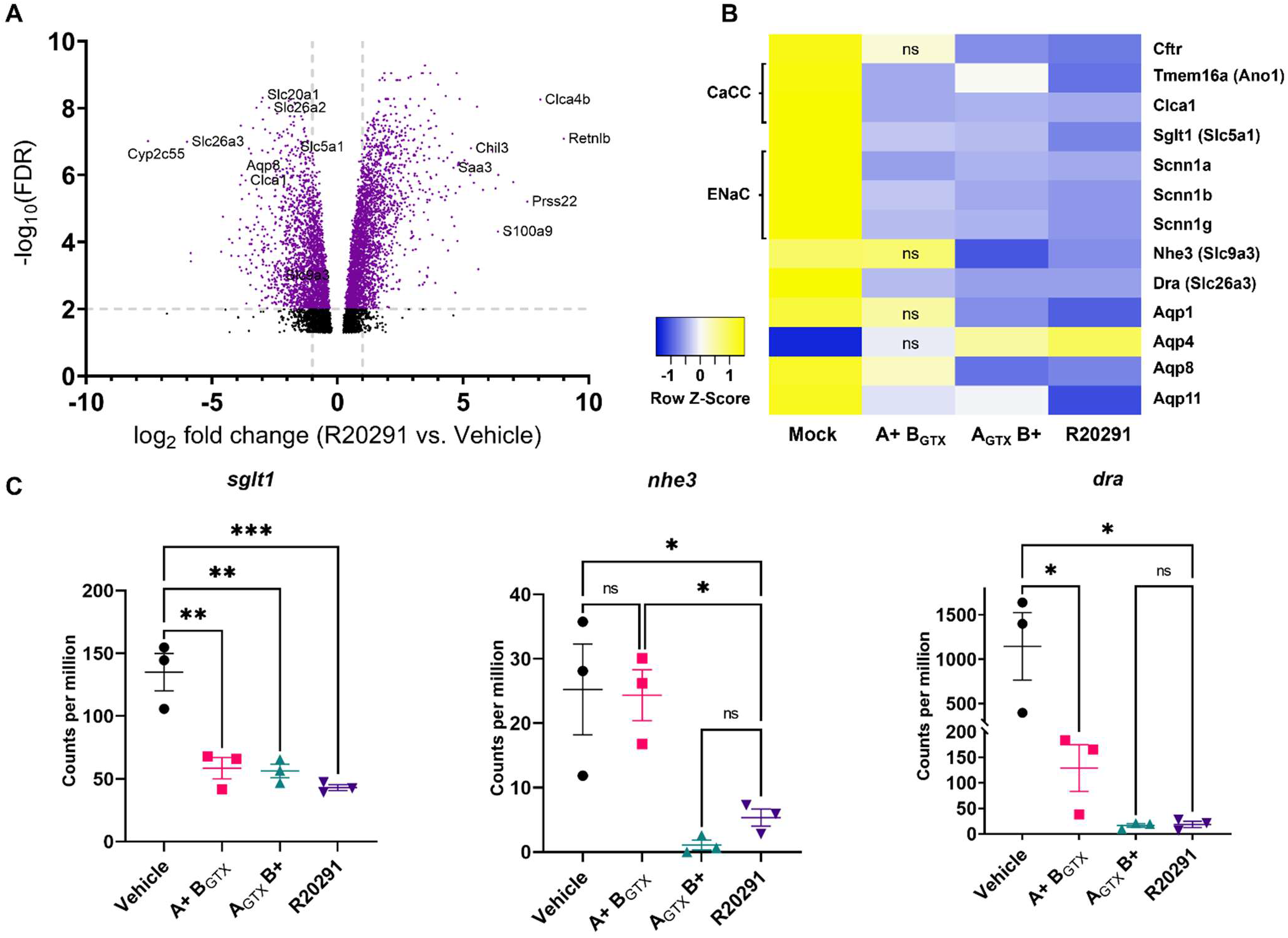
**A)** Volcano plot of differential expression data generated through RNA-sequencing of the distal colon during infection with R20291 (2 dpi) or in the vehicle control (PBS, 2 dpi). **B)** Expression heatmap of genes implicated in water and solute transport during infection. Expression of each gene is shown by *C. difficile* strain or vehicle control where high expression is shown in yellow and low in dark blue. **C)** Gene expression of *sglt1, nhe3,* and *dra* as determined by RNA-sequencing. Data are shown as counts per million, where the counts of each transcript were divided by the sum of library counts then multiplied by a million. * *P* < 0.05; ** *P* < 0.01; *** *P* < 0.001.

Transcript expression profile comparisons between vehicle control mice and A+ B_GTX_, A_GTX_ B+, and R20291-inoculated mice were conducted to explore toxin-dependent effects on expression (Supplemental Tables 2 and 3). Consistent with immunofluorescence protein expression data, there was a significant decrease in *Sglt1* expression in mice inoculated with mutant and wildtype strains (Figure 6B and 6C). Transcript data also supported TcdB-dependent repression of *Nhe3*, while TcdA had no effect on expression (Figure 6B and 6C). Finally, *Dra* expression was severely decreased during infection with mutant or wildtype strains, further supporting the sensitivity of its expression to both toxins (Figure 6B and 6C). These data imply that the absorptive ion transporters SGLT1, NHE3, and DRA are inhibited at the transcriptional level during CDI.

We assessed transcriptional changes in critical ion transporters that were tested using Üssing chambers. Unexpectedly, there were significant decreases in gene expression in *Cftr* when exposed to TcdB, but not TcdA, which may account for the numerical decrease in CFTR function during CDI that was statistically insignificant (Figures 3E and 6B). All three strains caused a significant decrease in the putative CaCCs, *Tmem16a (*also called *Ano1)* and *Clca1*. There was also significant repression of the ENaC-encoding genes *Scnn1a, Scnn1b,* and *Scnn1g* (Figure 6B). Decreases in transcript abundance of CaCC- and ENaC-encoding genes did not reflect the stable function of these ion transporter channels in the distal colon (Figures 3B, 3F and 6B), highlighting the complexity of expression and function of ion transporters during disease states.

Finally, aquaporins 1, 4, 8, and 11 were among the most differentially expressed genes during CDI (Figure 6B). Transcription of aquaporin 1 and 4-encoding genes was decreased or increased, respectively, dependent on TcdB (Figure 6B). TcdA could reduce expression of *Aqp8* and *Aqp11*, but *Aqp8* transcriptional repression was more potent in the presence of TcdB (Figure 6B). Decreased expression of *Aqp11* was most severe in R20291-infected mice, suggesting a synergistic effect in the presence of both TcdA and TcdB, which alone minimally reduced expression (Figure 6B). These data suggest that aquaporin proteins could also have a previously uncharacterized role in pathophysiology of water and solute transport during CDAD.

## DISCUSSION

Diarrhea during CDI can be mild and self-limiting, but in severe cases can cause dehydration through significant water and electrolyte loss. Additionally, it is the major source of pathogen spread between patients and onto environmental surfaces, where hardy spores can reside and become reservoirs for future outbreaks. Despite the importance of diarrhea during CDI, we do not fully understand the mechanisms underlying this symptom. Previous studies have used a mixture of cell culture models, intrarectal instillation of toxins in mice, ileal loop injections in rabbits, and human patient samples to examine the effects of toxins and CDI on expression of key ion transporters^33, 35, 36, 39, 40^. These studies implicated increased paracellular leakage and loss of expression of electroneutral ion exchangers NHE3, and DRA as potential causes of diarrhea. In this study, we sought to further define the factors underlying diarrhea and to determine the effect of *C. difficile* toxins on this process using a physiological infection model.

We had previously observed that diarrhea and weight loss severity were increased during infection with the epidemic B1/NAP1/PCR-ribotype 027 R20291 strain, when compared to mutants with impaired glucosyltransferase activity of TcdA or TcdB^28^. This led us to hypothesize that each toxin had unique effects on host physiology, that when synergized led to severe diarrhea. To begin to examine the role of *C. difficile* toxins on diarrhea, we first focused on defining regional changes in electrogenic ion transport in the ileum, cecum, proximal colon, and distal colon during infection with the wildtype R20291 strain.

Previous studies have tested the impact of toxins on permeability and Cl^-^ secretion using Üssing chambers^33, 57–59^. These studies utilized *ex vivo* rabbit ileal tissue that was exposed to crude or purified toxins inside the chamber system over the course of several hours. The murine CDI model allows us to study permeability and electrogenic ion transport in a physiological disease setting that includes a fully functioning immune response. We tested permeability *in vivo* by gavaging fasted mice with fluorescent dextran molecules of two different sizes. This experiment revealed that permeability is increased during CDI through the leak pathway, shown by increased 4 kDa FITC-dextran in plasma. Since *C. difficile* causes significant epithelial damage, we expected to see increased permeability of 70 kDa dextran through the unrestricted pathway, like what is observed in the dextran sodium sulfate model of colitis^47^. Based on previous studies, this result suggests that intestinal permeability during CDI is increased through tight junction-dependent mechanisms, specifically through the loss of occludin and phosphorylation of myosin light chain kinase 1^47, 60^. We tested tissue permeability in Üssing chambers to delineate where in the GI tract permeability was increased. Unexpectedly, there was no increase in tissue permeability in the ileum, cecum, proximal, or distal colon. This was a surprising result since there is extensive pathological damage in the cecum and colon during CDI. There are limitations to measuring tissue permeability in Üssing chambers, namely the small size of the tissue segment assayed. However, it is tempting to speculate that permeability may be increased in upper regions of GI tract (i.e., duodenum and/or jejunum) in response to pro-inflammatory signals generated in the cecum and colon during CDI.

Maintaining ion transport is essential for gut homeostasis and to balance solute absorption and secretion^61^. There were no significant changes in baseline transmucosal resistance (R_t_) or short circuit current in Üssing chambered tissues during CDI. Previous studies have found that both tissue R_t_ and transepithelial resistance in cell cultures are decreased after acute exposure to *C. difficile* toxins ^62–64^. To the best of our knowledge, this is the first time that R_t_ has been evaluated in tissues during a physiological *C. difficile* infection. These data along with permeability data suggest that there is a concerted response to maintain barrier function in terms of permeability during infection that may not be present in homogenous cell culture models and explant intoxication models. Indeed, studies have reported increased crypt length and epithelial cell differentiation during infection, which may reinforce the mucosal barrier during CDI^65, 66^. Chemical inhibition of ENaC revealed that there was no difference in function during infection. Contrary to findings from a previous study^64^, there was no increase in Cl^-^ secretion after addition of carbachol and forskolin. In fact, there was a decrease in Cl^-^ secretory potential in the cecum. There were no changes in CFTR function, and the loss of Cl^-^ secretion in the cecum was accounted for by reduced CaCC activity. The decrease in CaCC activity and Cl^-^ secretion may be a result of the severe epithelial damage that occurs in the cecum during murine CDI. Together, these results demonstrate that diarrhea during CDI is not a result of Cl^-^ hypersecretion, exemplified by chloride-rich cholera diarrhea, nor is it mediated by lack of sodium absorption by ENaC, which has been observed in Salmonella-induced enteritis^61, 67, 68^.

Üssing chamber experiments revealed that the function of SGLT1 was significantly impaired in the cecum and colon during CDI. Immunofluorescence imaging data further suggested that this loss of function is due to a near complete ablation of SGLT1 expression. We measured glucose concentration in the stool and there was a near two-fold increase in stool glucose during severe CDI compared to day zero. SGLT1-mediated carbohydrate absorption is thought to be mostly relevant in the small intestine and is only expressed at low levels in both rodent and human colons, thus, its function in the colon is incompletely understood^69, 70^. SGLT1 has been shown to actively uptake glucose in rat colons, albeit at a lower rate compared to the small intestine^71^. This result is intriguing to us since *in vitro* studies have shown that *C. difficile* represses toxin production in the presence of glucose through a conserved mechanism of carbon catabolite repression^72–74^. Moreover, a high carbohydrate diet rich in corn starch, casein, maltodextrin, and sucrose was protective against severe CDI outcomes^75^. Finally, the two dpi timepoint is when mice undergo the most severe CDI symptoms, and they typically begin resolving by 4 dpi as measured by weight gain and stool scores^28^. It is therefore conceivable that inactivation of SGLT1 in the colon during severe CDI would increase available glucose, decreasing toxin production, thus promoting disease resolution. Whether this is an intentional host-microbe interaction, or merely coincidence, remains unknown and merits further investigation.

We next sought to define the contributions of TcdA and TcdB on depletion of ion transporters implicated in diarrhea during CDI. Mice were inoculated with *C. difficile* mutants in the R20291 background with defined mutations that deactivate their glucosyltransferase activity^28^. We used immunofluorescence microscopy to image and quantify the relative abundance of SGLT1, NHE3, and DRA during acute infection. These experiments revealed that both TcdA and TcdB decrease DRA and SGLT1 expression significantly during infection, but only TcdB is capable of depleting NHE3. Unexpectedly, NHE3 expression was only decreased in the distal colon, where its expression is already very low, but not in the proximal colon. It has previously been shown that TcdB causes internalization of NHE3 in cell cultures^38^. The authors hypothesized that glucosylation of Rho GTPases by TcdB could lead to disruption of ERM and NHERF proteins, thus leading to internalization of NHE3^38^. We examined the expression of pERM and NHERF1 during infection and found no quantifiable differences. However, this does not preclude the possibility of internalization by this pathway, rather, we may be capturing a snapshot of infection where NHE3 has already been internalized and the cytoskeleton has undergone rearrangements that does not include returning NHE3 to the apical surface of colonocytes. This led us to hypothesize that these three ion transporters may be transcriptionally repressed.

We used an untargeted RNA-sequencing approach to further our understanding of dysregulation of ion transporters in the distal colon during infection. These data revealed a significant decrease in transcripts of *Sglt1, Nhe3, and Dra* during infection, and further supported TcdB-mediated dysregulation of NHE3 at the transcriptional level. This approach also highlighted four aquaporin proteins that were either up- or down-regulated during infection, which may play a role in water transport during diarrhea.

Transcriptional regulation of *Nhe3* is mediated by Sp1 and Sp3 transcription factors (specificity proteins 1 & 3), and transcription of *Sglt1* is regulated by Sp1 and HNF1 (hepatocyte nuclear factor 1)^76, 77^. Gene expression of *Dra* is regulated by direct binding of CDX2 and NF-κB (caudal-type homeobox protein 2 & nuclear factor kappa-light-chain-enhancer of activated B cells) to different sites in the *Dra* promoter^78, 79^. These transcription factors are subject to several regulatory stimuli including butyrate and the proinflammatory cytokines TNFα (tumor necrosis factor-alpha), IFNγ (interferon gamma), and IL-1β (interleukin 1 beta)^79, 80^. Sp1 and Sp3 are inhibited by PKA-mediated phosphorylation induced by TNFα^76^, HNF1 and CDX2 are transcriptionally repressed by TNFα^81, 82^, and NF-κB is potently activated by TNFα^83^. Additionally, paracellular flux through the leak pathway is remarkably upregulated in response to T cell-mediated increases in TNFα and IFNγ^60, 84^. The combination of these findings lead us to hypothesize that upregulation of proinflammatory cytokines increases paracellular flux of water and solutes in upper areas of the small intestine through the leak pathway, and decrease solute absorption by NHE3, DRA, and SGLT1 in the colon, leading to coordinated dysfunction that causes osmotic diarrhea during CDI. Further experiments are needed to define the inflammatory pathways that underlie these pathophysiological changes in solute and water transport.

In conclusion, these data demonstrate that both TcdA and TcdB are necessary in the RT 027 R20291 strain to cause the starkest pathophysiological changes that lead to diarrhea during CDI. There was an increase in paracellular permeability through the leak pathway during infection, but potentially not in the lower GI where CDI symptoms and pathology are more pronounced. Additionally, there was little change in transmucosal resistance or short circuit current in the ileum, cecum, proximal colon, and distal colon when tested in Üssing chambers. In fact, the only changes in electrogenic ion transporters measured were decreased CaCC activity in the cecum and loss of SGLT1 function in the cecum, proximal colon, and distal colon. Further experiments demonstrated a role for either or both TcdA and TcdB in reducing SGLT1, NHE3, and DRA at the protein and mRNA levels. Questions remain about the pathophysiological cues that are involved in dysregulation of these transporters, which will be addressed in future studies.

## Abbreviations used in this paper

CDI: Clostridioides difficile infection

GI: gastrointestinal

NHE3: Na+/H+ Exchanger 3

DRA: Downregulated in Adenoma

BHIS: brain-heart infusion-supplemented medium

TA: taurocholic acid

TCCFA: taurocholic acid, D-cycloserine, cefoxitin and fructose agar

FITC: fluorescein isothiocyanate

FD4: 4 kDa FITC dextran

RD70: 70 kDa rhodamine B dextran

KRB: Krebs-Ringer Buffer

Isc: short-circuit current

Rt: transmucosal resistance

Gt: tissue conductance

SGLT1: sodium-glucose cotransporter 1

ENaC: epithelial sodium channel

CFTR: cystic fibrosis transmembrane conductance regulator

CaCC: calcium activated chloride channels

PFA: paraformaldehyde

NBF: neutral buffered formalin

FFPE: formalin-fixed paraffin embedded

OCT: optimal cutting temperature embedding medium

ERM: Ezrin/Radixin/Moesin

NHERF: Na+/H+ exchanger regulatory factor

Sp1 & Sp3: specificity proteins 1 and 3

HNF1: hepatocyte nuclear factor 1

CDX2: caudal-type homeobox protein 2

NF-κB: nuclear factor kappa-light-chain-enhancer of activated B cells

TNFα: tumor necrosis factor-alpha

IFNγ: interferon gamma

IL-1β: interleukin 1 beta

BI/NAP1/PCR-RT: restriction endonuclease analysis type B1, North American pulse-field gel electrophoresis type 1 polymerase chain reaction ribotype

## ACKNOWLEDGEMENTS

Thank you to the members of the Lacy lab for stimulating discussions and feedback on this manuscript. Thank you Kari Seedle and the Vanderbilt Cell Imaging Shared Resource Core for support on imaging and analyses; Vanderbilt Technologies for Advanced Genomics for support with RNA-sequencing; and TPSR for histology support. The graphical abstract was made using BioRender.

## TABLE KEY RESOURCES

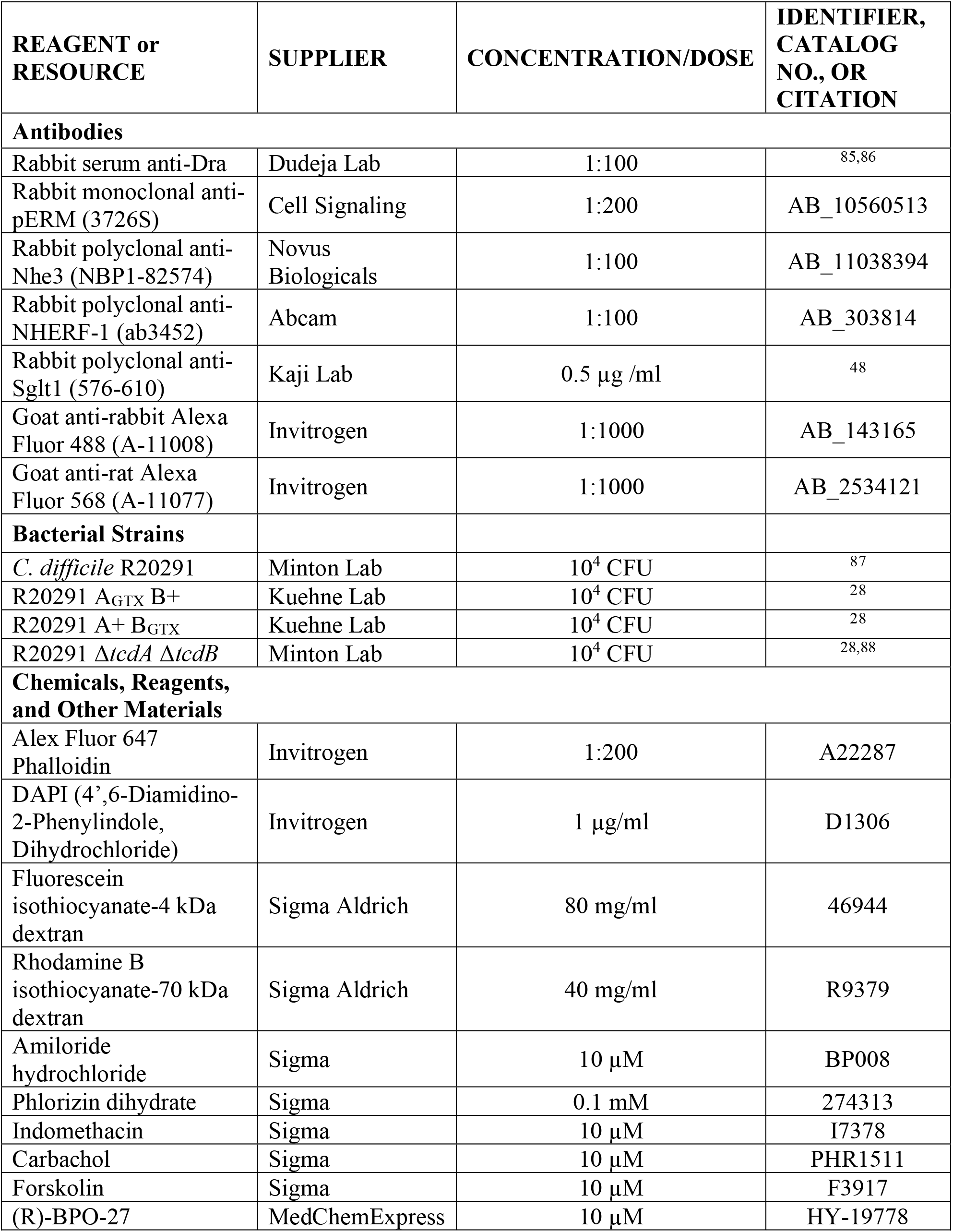
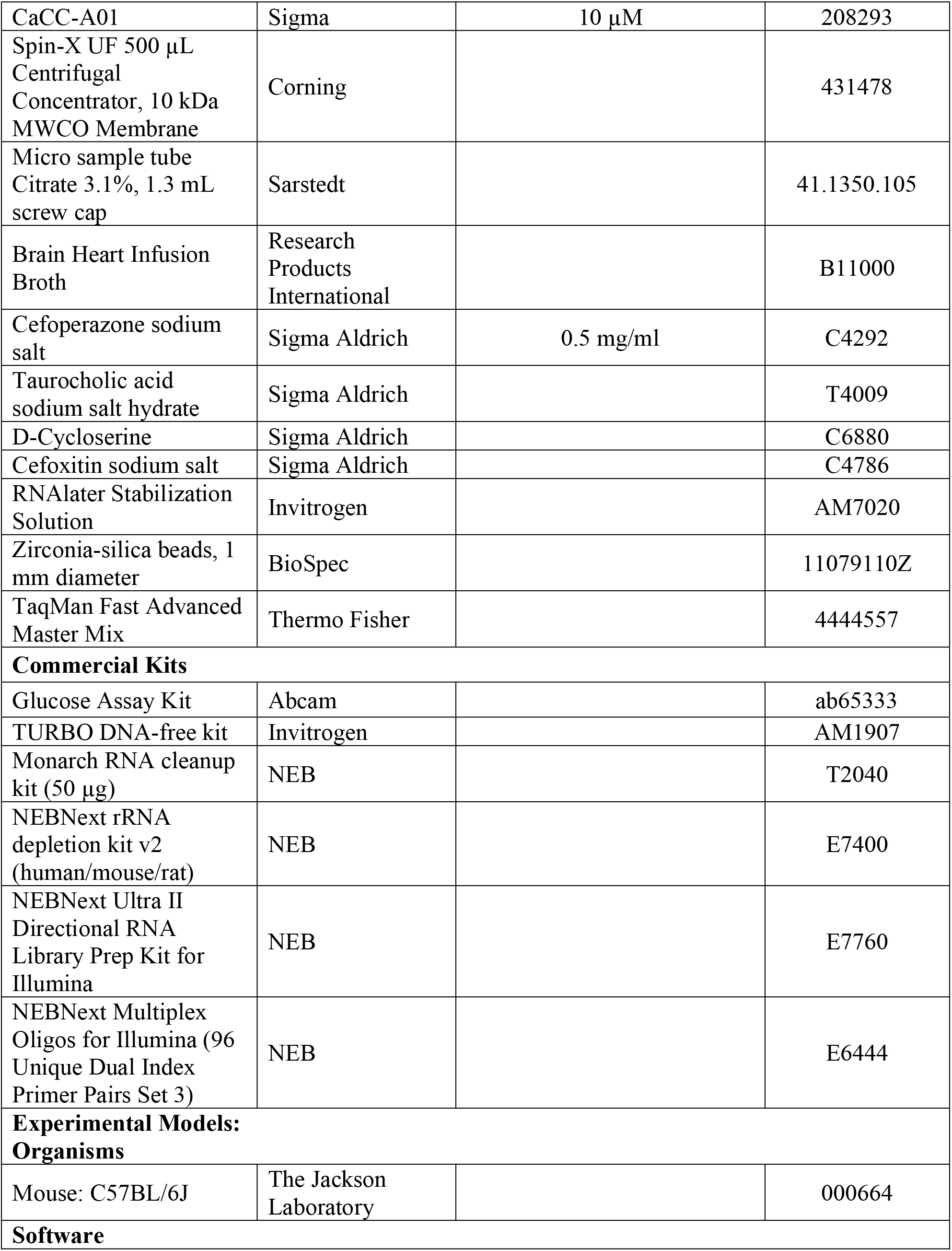
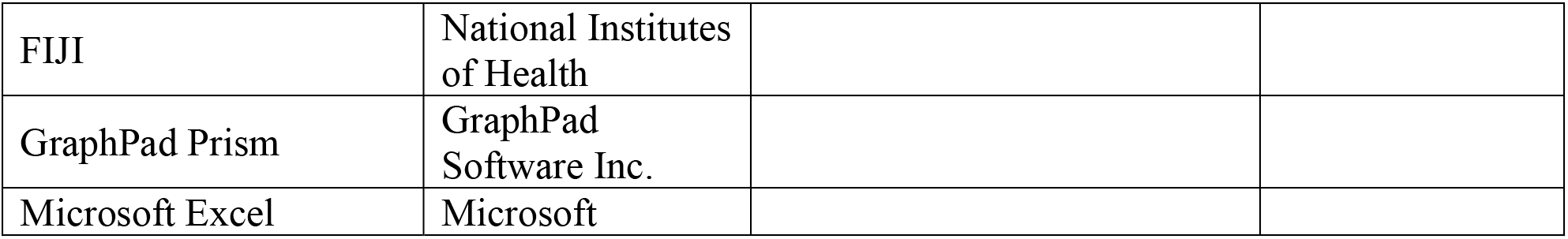

| REAGENT or RESOURCE | SUPPLIER | CONCENTRATION/DOSE | IDENTIFIER, CATALOG NO., OR CITATION |
| --- | --- | --- | --- |
| **Antibodies** |  |  |  |
| Rabbit serum anti-Dra | Dudeja Lab | 1:100 | 85,86 |
| Rabbit monoclonal anti-pERM (3726S) | Cell Signaling | 1:200 | AB_10560513 |
| Rabbit polyclonal anti-Nhe3 (NBP1-82574) | Novus Biologicals | 1:100 | AB_11038394 |
| Rabbit polyclonal anti-NHERF-1 (ab3452) | Abcam | 1:100 | AB_303814 |
| Rabbit polyclonal anti-Sglt1 (576-610) | Kaji Lab | 0.5 µg /ml | 48 |
| Goat anti-rabbit Alexa Fluor 488 (A-11008) | Invitrogen | 1:1000 | AB_143165 |
| Goat anti-rat Alexa Fluor 568 (A-11077) | Invitrogen | 1:1000 | AB_2534121 |
| **Bacterial Strains** |  |  |  |
| *C. difficile* R20291 | Minton Lab | 10<sup>4</sup> CFU | 87 |
| R20291 A<sub>GTX</sub> B<sup>+</sup> | Kuehne Lab | 10<sup>4</sup> CFU | 28 |
| R20291 A<sup>+</sup> B<sub>GTX</sub> | Kuehne Lab | 10<sup>4</sup> CFU | 28 |
| R20291 $\Delta tcdA \Delta tcdB$ | Minton Lab | 10<sup>4</sup> CFU | 28,88 |
| **Chemicals, Reagents, and Other Materials** |  |  |  |
| Alex Fluor 647 Phalloidin | Invitrogen | 1:200 | A22287 |
| DAPI (4',6-Diamidino-2-Phenylindole, Dihydrochloride) | Invitrogen | 1 µg/ml | D1306 |
| Fluorescein isothiocyanate-4 kDa dextran | Sigma Aldrich | 80 mg/ml | 46944 |
| Rhodamine B isothiocyanate-70 kDa dextran | Sigma Aldrich | 40 mg/ml | R9379 |
| Amiloride hydrochloride | Sigma | 10 µM | BP008 |
| Phlorizin dihydrate | Sigma | 0.1 mM | 274313 |
| Indomethacin | Sigma | 10 µM | I7378 |
| Carbachol | Sigma | 10 µM | PHR1511 |
| Forskolin | Sigma | 10 µM | F3917 |
| (R)-BPO-27 | MedChemExpress | 10 µM | HY-19778 |
| CaCC-A01 | Sigma | 10 μM | 208293 |
| Spin-X UF 500 μL Centrifugal Concentrator, 10 kDa MWCO Membrane | Corning | | 431478 |
| Micro sample tube Citrate 3.1%, 1.3 mL screw cap | Sarstedt |  | 41.1350.105 |
| Brain Heart Infusion Broth | Research Products International |  | B11000 |
| Cefoperazone sodium salt | Sigma Aldrich | 0.5 mg/ml | C4292 |
| Taurocholic acid sodium salt hydrate | Sigma Aldrich |  | T4009 |
| D-Cycloserine | Sigma Aldrich |  | C6880 |
| Cefoxitin sodium salt | Sigma Aldrich |  | C4786 |
| RNAlater Stabilization Solution | Invitrogen |  | AM7020 |
| Zirconia-silica beads, 1 mm diameter | BioSpec |  | 11079110Z |
| TaqMan Fast Advanced Master Mix | Thermo Fisher |  | 4444557 |
| **Commercial Kits** |  |  |  |
| Glucose Assay Kit | Abcam |  | ab65333 |
| TURBO DNA-free kit | Invitrogen |  | AM1907 |
| Monarch RNA cleanup kit (50 μg) | NEB | | T2040 |
| NEBNext rRNA depletion kit v2 (human/mouse/rat) | NEB |  | E7400 |
| NEBNext Ultra II Directional RNA Library Prep Kit for Illumina | NEB |  | E7760 |
| NEBNext Multiplex Oligos for Illumina (96 Unique Dual Index Primer Pairs Set 3) | NEB |  | E6444 |
| **Experimental Models: Organisms** |  |  |  |
| Mouse: C57BL/6J | The Jackson Laboratory |  | 000664 |
| **Software** |  |  |  |
| FIJI | National Institutes<br>of Health |  |  |
| GraphPad Prism | GraphPad<br>Software Inc. |  |  |
| Microsoft Excel | Microsoft |  |  |

---

**SUPPLEMENTARY FIGURE S1.**
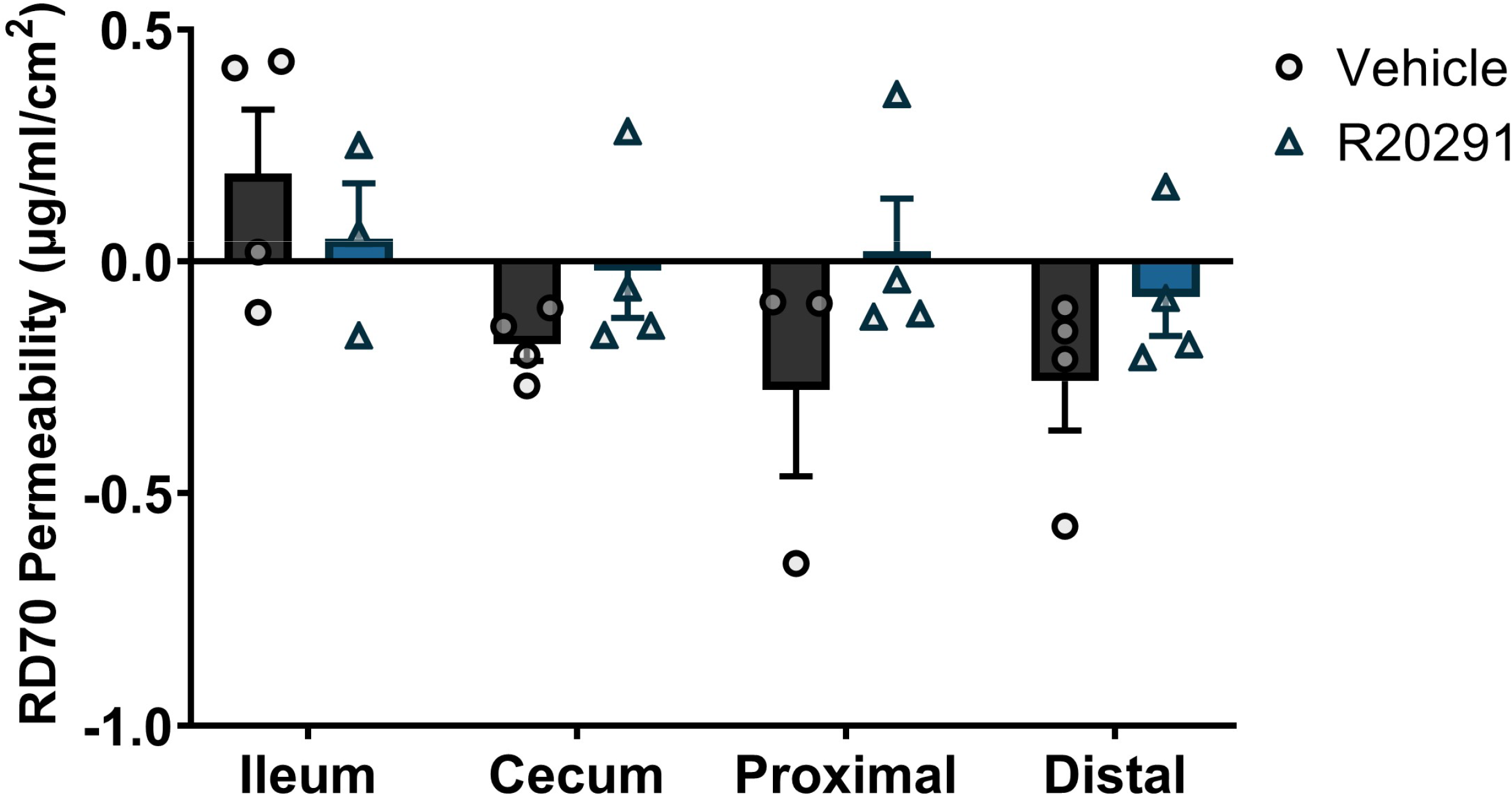
There was no difference in paracellular permeability between mock- and R20291-inoculated mice at 2 dpi via the unrestricted pathway in the ileum, cecum, proximal colon, and distal colon (*n* = 3-4 per treatment per region).

**SUPPLEMENTARY FIGURE S2.**
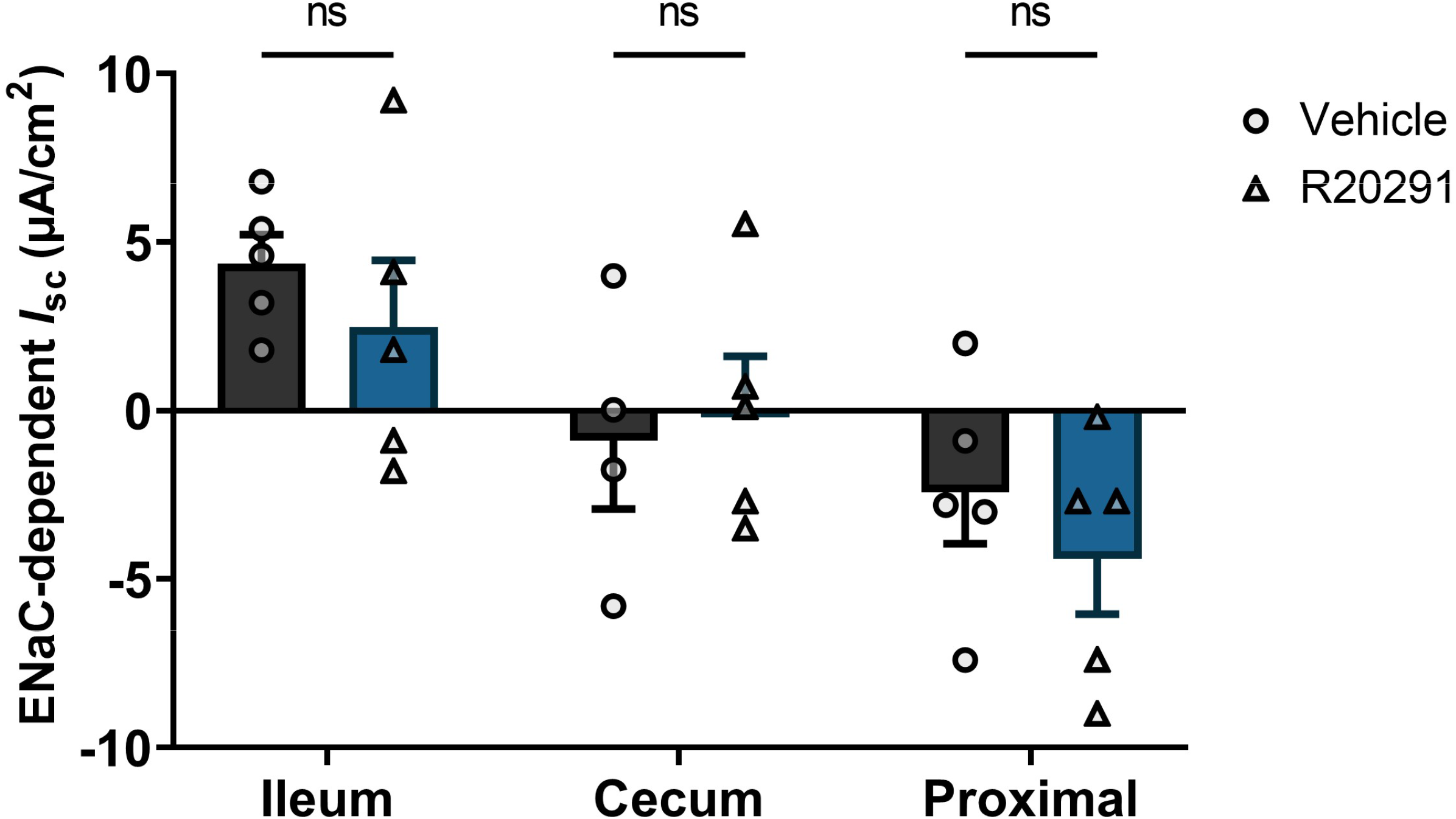
The absorptive sodium channel, ENaC, is mostly functionally active in the distal colon. Its function is unchanged in the ileum, cecum, and proximal colon during CDI.

**SUPPLEMENTARY FIGURE S3.**
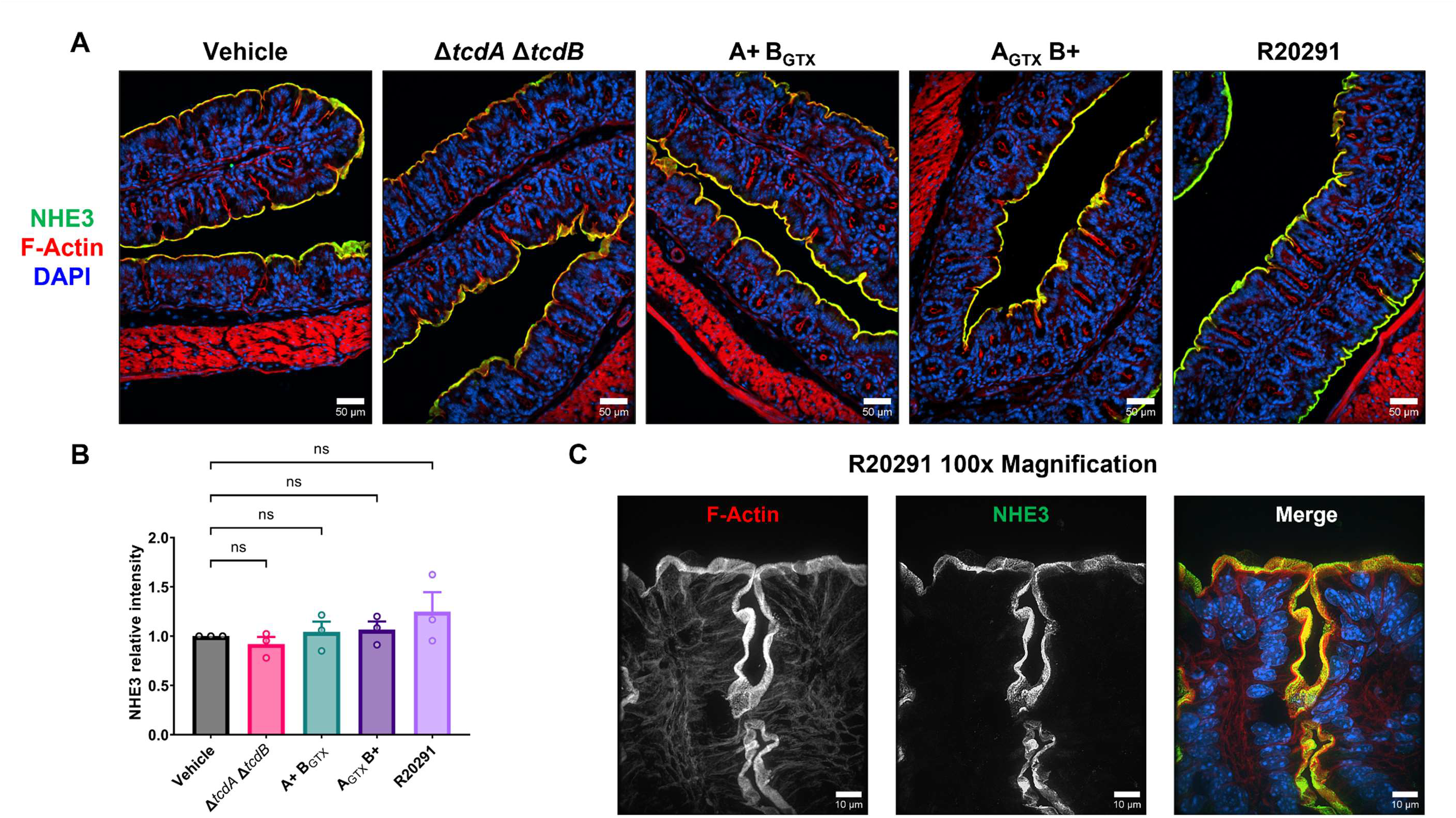
**A & B)** Immunofluorescence imaging (20x magnification) of NHE3 demonstrates that expression is unchanged in the proximal colon during CDI when infected with mutant and wildtype *C. difficile* strains (2 dpi). Scale bars = 50 µm **C)** Higher magnification provides further insight into the sustained apical localization of NHE3 in the proximal colon of *C. difficile* infected mice. Scale bars = 10 µm.

**SUPPLEMENTARY FIGURE S4.**
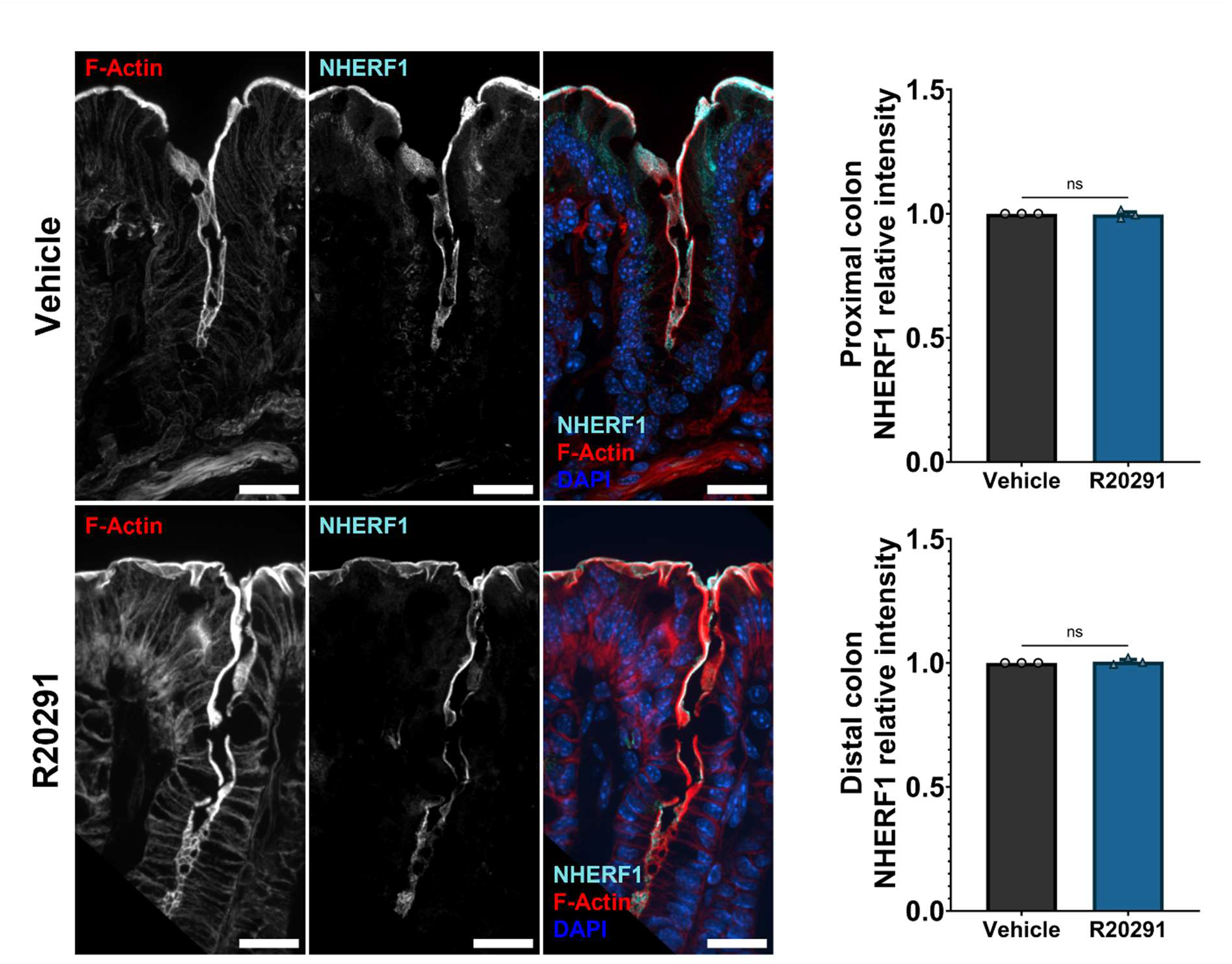
Immunofluorescence imaging of NHERF1 shows that expression is unchanged in both proximal and distal colon regions during CDI. Images are 60x magnification and scale bars represent 20 µm.

**SUPPLEMENTARY FIGURE S5.**
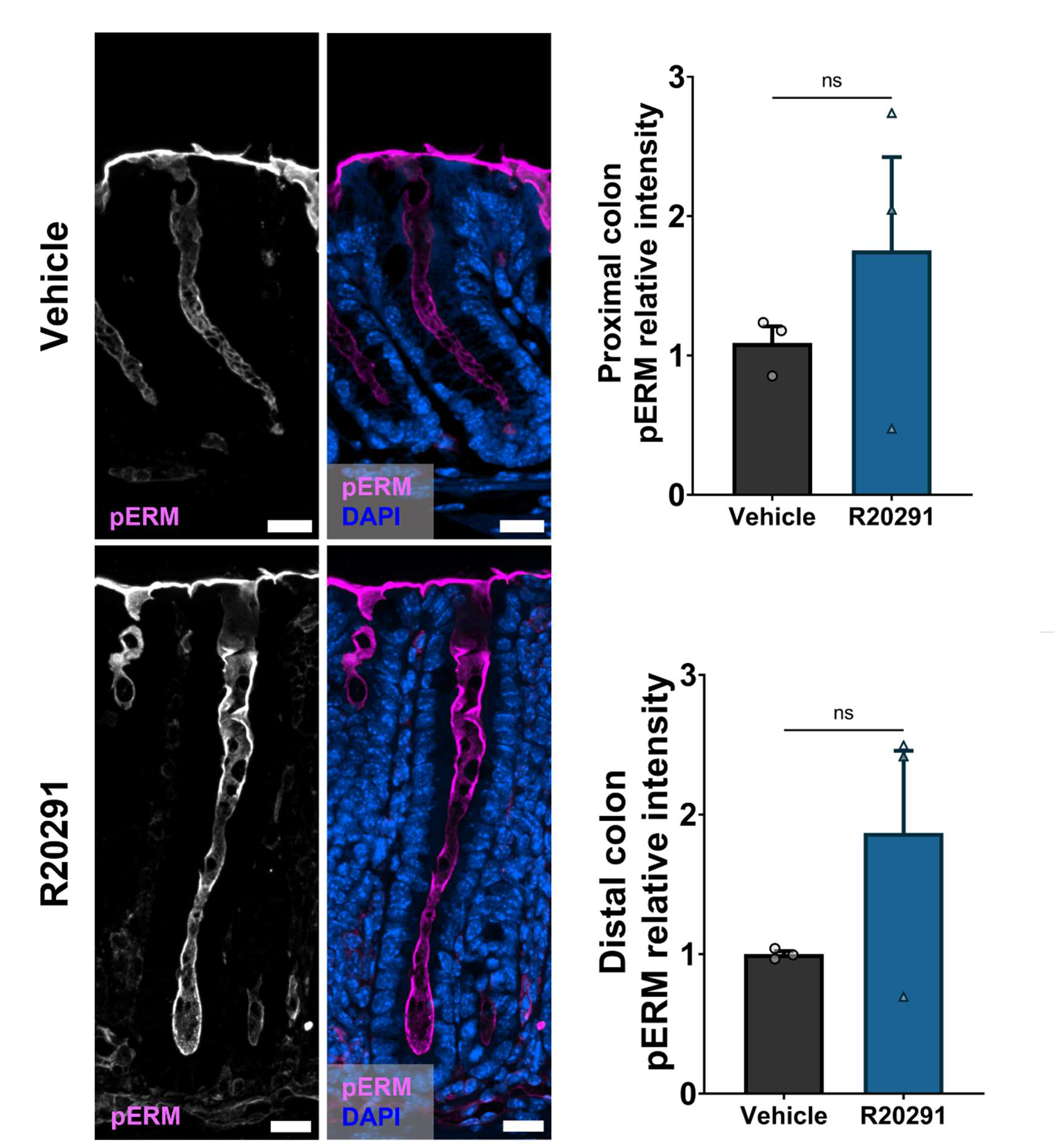
Immunofluorescence imaging of pERM demonstrates that expression is insignificantly unchanged in both proximal and distal colon regions during CDI. Images are 40x magnification and scale bars are 20 µm.

